# Clonal sharing of CD8+ T-cells links skin and joint inflammation in psoriatic arthritis

**DOI:** 10.1101/2024.05.09.593313

**Authors:** Lucy E. Durham, Frances Humby, Nora Ng, Elizabeth H. Gray, Sarah E. Ryan, Rosie Ross, Giovanni A. M. Povoleri, Rosamond Nuamah, Kathy Fung, Athul Menon Kallayil, Pawan Dhami, Bruce W. Kirkham, Leonie S. Taams

## Abstract

We hypothesised that skin and joint inflammation in psoriatic arthritis (PsA) is linked in terms of CD8+ T-cell phenotype and clonality. We employed scRNAseq to directly compare the transcriptional signature and T-cell receptor repertoire of memory T-cells from paired skin and synovial tissue and/or fluid from patients with PsA. We identified an enrichment of type-17 CD8+ tissue-resident memory (T_RM_) T-cells in both skin and joint, with a stronger IL-17 signature in the skin than the joint. Several T-cell clones were shared between the skin and joint and these shared clones tended to have the same signature at both sites, characterised by increased expression of genes associated with a cytotoxic, tissue-resident phenotype. Our findings support the hypothesis that skin and joint inflammation in PsA is linked in terms of T-cell clonality and raises the possibility that specific T-cells migrate between these compartments to propagate inflammation across both sites.

## Introduction

Psoriatic arthritis (PsA) is an inflammatory arthritis that develops in up to 30% of people with the skin condition psoriasis^1^. It is unclear what initiates or drives chronic inflammation in PsA or if the immune mechanisms underlying skin and joint inflammation are the same. We hypothesised that skin and joint inflammation in PsA is linked in terms of CD8+ T-cell gene signature and clonality.

Evidence strongly supports a role for both IL-17 and antigen-driven activation of CD8+ T cells in inflammation in psoriasis and PsA. Type-17, tissue resident CD8+ T-cells are enriched in both the skin in psoriasis and in the joints in PsA pointing to a key role for these cells in driving inflammation in both sites^2–8^. Furthermore, monoclonal antibodies that target IL-17A/F treat both psoriasis and PsA confirming the clinical relevance of the IL-17 pathway in both diseases^9,10^. As well as pointing to a role for the IL-17 pathway, the genetics of both psoriasis and PsA also support a role for MHC class I-mediated presentation of antigen to CD8+ T cells^11–15^. Both diseases are associated with specific HLA class I alleles and with single nucleotide polymorphisms in *ERAP1*/2, required for HLA class I peptide trimming, and *RUNX3,* which is essential for CD8+ T-cell differentiation^11–15^.

The T-cell receptor (TCR) repertoire in inflamed joints is distinct from that in blood, suggesting antigen-driven recruitment and/or activation of synovial T-cells in PsA^16^. Furthermore, one study reported the presence of identical TCRβ sequences in paired skin and synovium in PsA, suggesting that a common antigen may be present in both sites^17^. However, this study only sequenced a small number of TCRβ sequences and did not investigate the phenotype of shared clones^17^. In both psoriasis and PsA, effective treatment of inflammation is associated with dispersion of the polyclonal T-cell infiltrate leaving a comparatively oligoclonal population, which in the skin has been shown to express IL-17^18,19^. This resident population of clonally expanded, IL-17+ T-cells is thought to be the critical driver of disease recurrence upon cessation of treatment^3,19,20^.

Despite compelling evidence for a link between PsA and psoriasis, differences have been noted, including different HLA class I gene associations and risk alleles identified by Genome Wide Association Studies (GWAS)^11,21,22^. Clinically, the two diseases can have different therapy responses, particularly to ciclosporin and sulfasalazine^23,24^. Lastly, in rare studies where paired skin and synovial tissue (ST) were directly compared, gene expression profiling revealed a stronger IL-17/IL-23 signature in the skin compared to TNFα, IL-6 and IFNγ signatures in ST^25,26^. In these studies, RNA was extracted from whole tissue and therefore gene expression reflected a pool of different immune and stromal cells, which may explain disease characteristics but does not address key pathways driving psoriatic disease. The limitation of the few studies to date that have directly compared skin and synovium in PsA is that they have compared *either* the TCR sequence *or* the phenotype of bulk cells. To our knowledge, no study has simultaneously compared the phenotype *and* TCR sequence of single T-cells. To address this knowledge gap, we used single cell RNA sequencing (scRNAseq) to directly compare the gene signature and TCR repertoire of memory T-cells from paired skin and ST from patients with PsA. We demonstrate an enrichment of type-17 CD8+ tissue-resident memory (T_RM_) cells in both skin and joint compared to blood, with a stronger IL-17 signature in the skin. We also identify several CD8+ T-cell clones, that are shared between the skin and the joint and have similar signatures at both sites, characterised by increased expression of genes associated with a cytotoxic, tissue-resident phenotype. This supports the hypothesis that skin and synovial inflammation is linked in terms of T-cell clonality and raises the possibility that specific T-cells migrate between the skin and the joint to propagate inflammation across both sites.

## Results

Six patients with PsA donated paired samples of inflamed skin, ST and/or synovial fluid (SF) from an inflamed knee joint and blood (**Table 1** details demographic and clinical information). HLA serotyping was available for 5 of the 6 patients (**Table 1**). Of note, PsA 1 and PsA 5 both expressed the major risk allele for psoriasis, *HLA-C*06:02*; PsA 1 and PsA 3 expressed the haplotype *HLA-B*08:01-HLA-C*07:01*, which is associated with severe PsA, and PsA 4 and PsA 6 both expressed *HLA-B*38:01-HLA-C*12:03*, which has also been associated with PsA^11,27^.

**Table 1.**
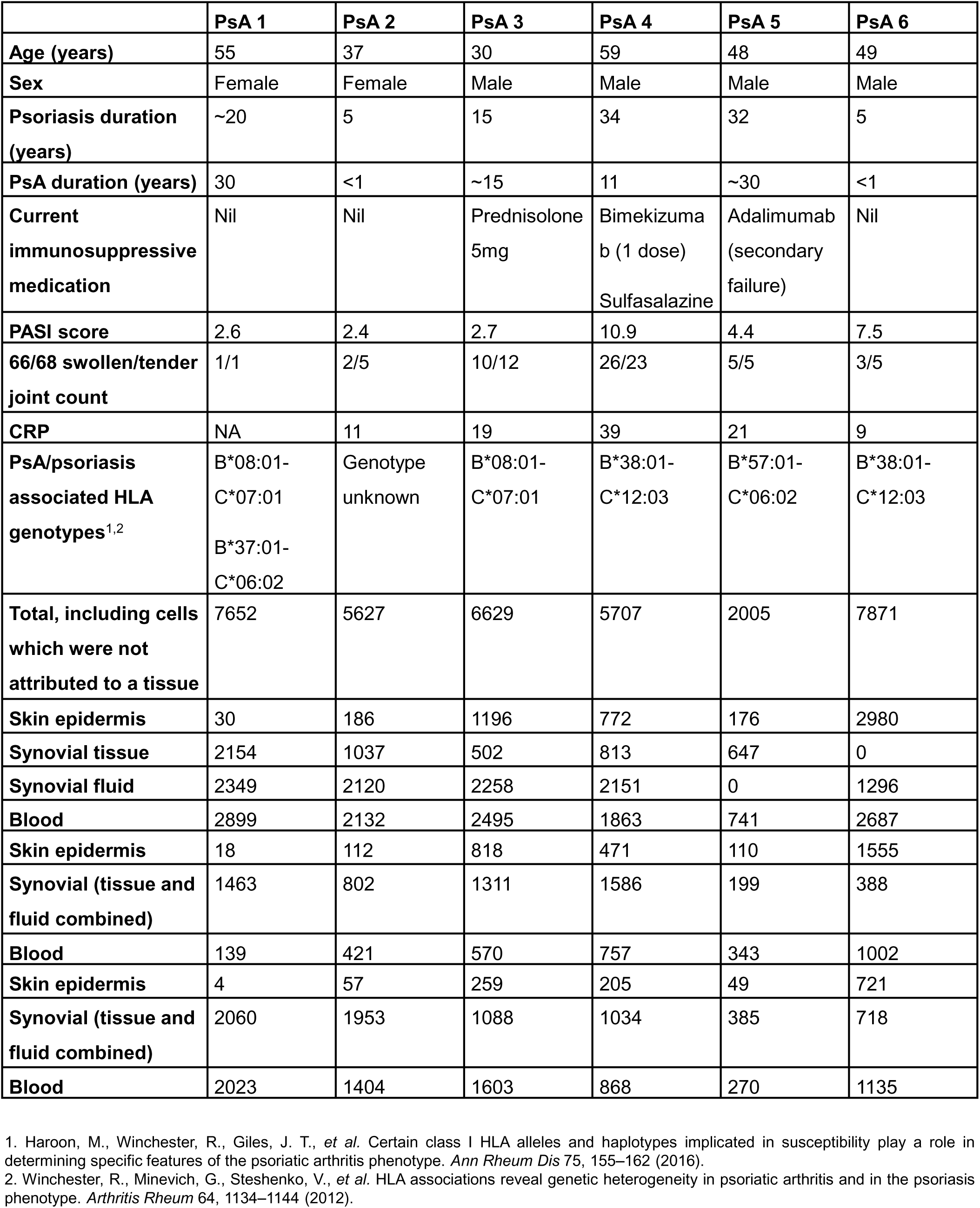
Participant information, psoriasis/PsA associated HLA genotypes, and single cell yield from each participant.

Live memory (CD45RA-CD27+/- and CD45RA+CD27-) T-cells were sorted from each sample (**Figure S1)** for scRNAseq using the 10x Genomics platform. To eliminate within-patient batch effects arising from library preparation or sequencing, cells from each tissue were barcoded with CITE-seq hashtag antibodies and combined prior to loading on to the 10x Genomics chip. Following demultiplexing, QC (**Figure S2**), and normalisation with regression of the effects of cell cycle genes, the percentage of mitochondrial genes and digestion module score^28^, 35,491 memory T-cells from the six patients were integrated and clustered in Seurat (**Figure S3, Table 1** details cell yield by patient)^29–31^. Memory CD8+ and CD4+ T-cell subsets were then identified and analysed independently^28^ (**Figure S3**).

### Enrichment of CD8+ T_RM_ cells in skin and joint compared to blood

Seurat clustering of 15,335 memory CD8+ T-cells from blood, skin epidermis, ST and/or SF from the six patients yielded 19 clusters (**Figure 1a, Supplementary Data 1**). Whilst skin and blood CD8+ T-cells clustered distinctly from each other on the UMAP, ST and SF T-cells had overlapping clustering profiles (indicating similarity) and were spread more diffusely across the whole UMAP indicating that some synovial cells had a gene signature similar to cells from blood, some to those from skin and others (for example those residing in clusters 0, 8, and 10) were more specific to the joint (**Figure 1b, c**).

**Figure 1:**
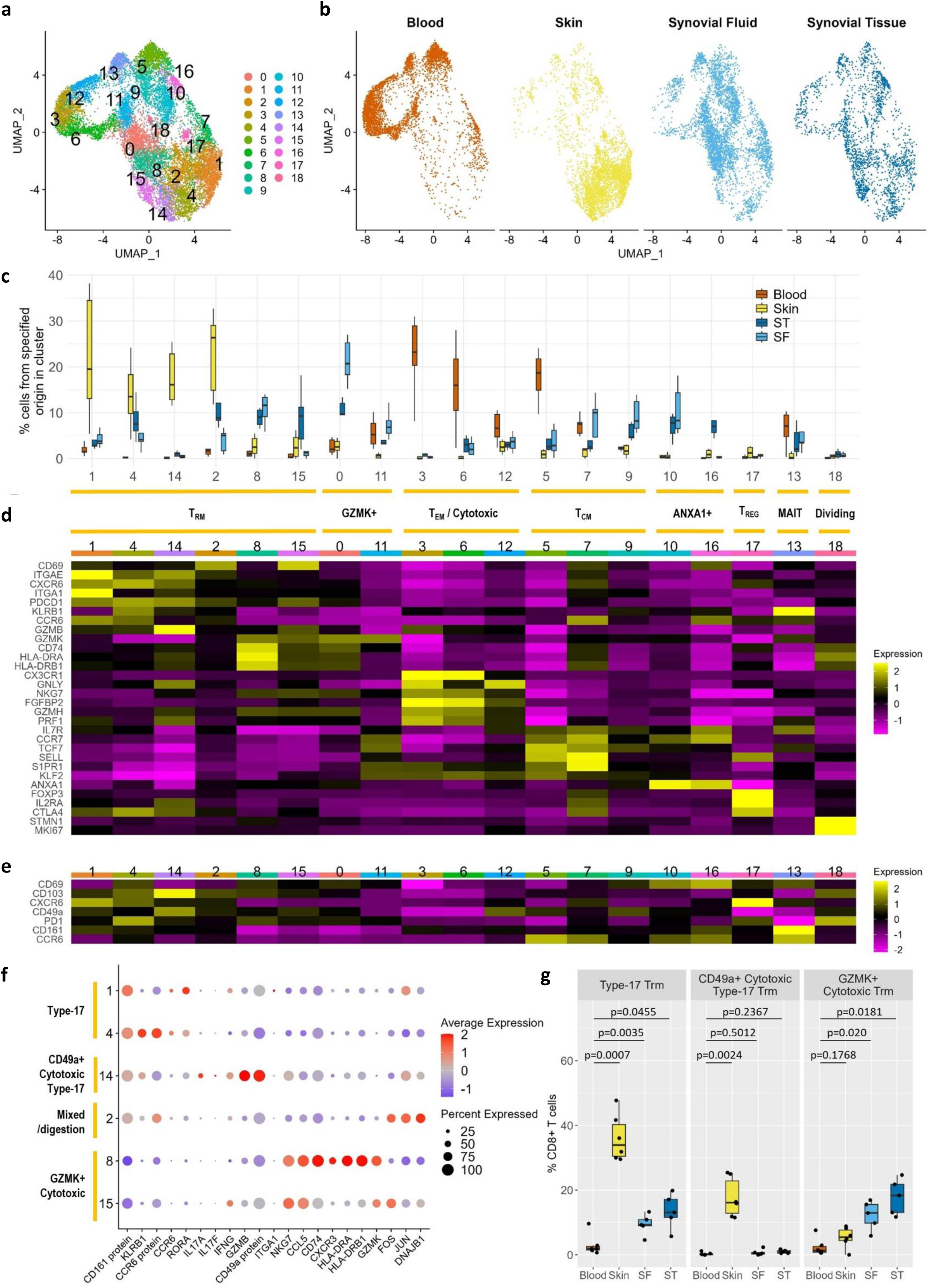
Signature of memory CD8+ T-cells in PsA. **a)** UMAPs of integrated analysis of 15,335 memory CD8+ T-cells from paired samples of blood, lesional skin epidermis and synovial tissue and/or synovial fluid from an inflamed joint (n=6 patients). Each dot represents a cell. Proximity of dots to each other indicates the similarity of the cells. Cells coloured according to the 19 cell populations obtained after Seurat clustering. **b)** UMAPs split by tissue of origin. **c)** Distribution of CD8+ T-cells from each tissue across the 19 clusters (n=6 patients). Clusters are ordered in groups with similar phenotypes. **d,e)** Heatmaps visualising averaged expression of specified genes **(d)** and proteins **(e)** in each cluster. **f)** Dot plot depicting the expression of select genes across the CD8+ T_RM_ clusters. Size of dots indicates the % of cells within the cluster that express the indicated gene. Colour of dots indicates the scaled expression of indicated gene across the CD8+ T_RM_ clusters. Yellow bars indicate clusters with similar phenotypes. **g)** Frequency of total memory CD8+ T-cells from blood, skin, SF, ST within each indicated T_RM_ subset. Mixed-effects analysis comparing skin (n=6) to blood (n=6), SF (n=5) to blood (n=6), and ST (n=5) to blood (n=6)) with Dunnett’s correction for multiple comparisons. Adjusted p values shown.

Analysis of differentially expressed genes between clusters (**Figure S4, Supplementary Data 2**) enabled grouping of clusters with similar signatures (**Figure 1c, d, e**). The major CD8+ T-cell subsets known to be present in inflamed tissues in psoriasis and PsA were identified including T_RM_ cells (clusters 1, 2, 4, 8, 14 and 15), granzyme K+ (clusters 0 and 11), cytotoxic (clusters 3, 6 and 12), central memory (clusters 5, 7 and 9), Annexin A1 (ANXA1)+ (clusters 10 and 16), T_REG_-like (cluster 17), mucosal-associated invariant T (MAIT) (cluster 13 which had significantly upregulated expression of *TRAV1-2* and *KLRB1*, **Supplementary Data 2**), and dividing (cluster 18) cells^4,7,8,16,32^.

Since T_RM_ cells are enriched at sites of inflammation in both psoriasis and PsA ^3,4,6–8^ and are hypothesised to contribute to the persistence of both diseases^3,7,19,33^, these cells were investigated in more detail. To identify T_RM_ cell clusters, we utilised differential gene expression in combination with GSEA^28^. Under homeostatic conditions, human T_RM_ cells are frequently identified as CD69+CD103+/-^34^. However, in our dataset, CD69 RNA/protein was diffusely expressed across most clusters which contained skin and synovial cells, and likely reflected T-cell activation rather than residency (**Figure S5**). Hence, CD69 expression in isolation, was not sufficient to identify CD69+CD103- T_RM_ cells in this context. Therefore, we defined “potential T_RM_” clusters (clusters 0, 1, 2, 4, 8, 14 and 15) as clusters with significant differential expression of 3+ genes/proteins associated with the “core-transcriptional signature” of human T_RM_ cells (increased CD103/*ITGAE*, CD69/*CD69*, CXCR6/*CXCR6*, CD49a/*ITGA1*, PD-1/*PDCD1*, decreased *S1PR1, SELL, KLF2,* with at least one gene/protein increased and one decreased)^34^ (**Figure S6a, Supplementary Data 2**). GSEA was then performed for each potential T_RM_ cluster compared to pooled cells from “non-potential-T_RM_” clusters. Clusters that were positively enriched for a core T_RM_ signature from homeostatic lung^34^ (clusters 1, 2, 4, 8, 14 and 15) were classified as T_RM_ clusters (**Figure S6b)**.

Further examination of gene expression (**Figure 1f, Supplementary Data 2**) and GSEA comparing each T_RM_ cluster to pooled cells from the other T_RM_ clusters (**Figure S6c**) identified three key CD8+ T_RM_ cell subsets. Type-17 T_RM_ cells (clusters 1 and 4), which were significantly enriched in the skin and also in the joint (as we have previously shown^6^) compared to the blood; CD49a+ cytotoxic type-17 T_RM_ cells (cluster 14), which were significantly enriched in the skin compared to the blood, and GZMK+ cytotoxic T_RM_ cells (clusters 8 and 15), which were significantly enriched in the joint compared to the blood (**Figure 1g)**. T_RM_ cell cluster 2 was not enriched for either a type-17 or a CD49a+GZMK+ T_RM_ signature but was positively enriched for a signature associated with tissue that has undergone collagenase digestion^35^ and was therefore classified as a “mixed/digestion” signature subset **(Figure 1f)**.

Cluster 0 met the criteria to be a “potential-T_RM_ cluster” but was not positively enriched for a T_RM_ signature on GSEA (**Figure S6b**). It contained predominantly synovial cells (**Figure 1c**) that expressed high levels of GZMK and HLA-DR genes (**Figure 1d, Supplementary Data 2**), similar to SF CD49a+GZMK+ T_RM_ cells we previously reported in PsA^7^. Cluster 0 shared high expression of *GZMK* with cluster 11 and therefore was classified as “GZMK+” (**Figure 1d**). We speculate that cells in cluster 0 may be transitioning from a GZMK+ phenotype shared with blood (cluster 11) towards a GZMK+ T_RM_ phenotype (clusters 8 and 15).

### Striking differences in the phenotype of skin vs synovial CD8+ T-cells

Having identified the expected CD8+ T-cell subsets in the dataset, we sought to directly compare memory CD8+ T-cells from skin epidermis with ST. Comparison of gene expression revealed that skin CD8+ T-cells had significantly increased expression of *CXCR6, IFNG* and *GZMB* and a stronger IL-17 signature with significantly increased expression of *IL17A* and *CCL20* compared to ST (**Figure 2a, Supplementary Data 3**), and increased expression of other type-17 genes including *IL17F, IL26* and *IL23R* (**Figure 2b**). In contrast, ST CD8+ T-cells had significantly increased expression of *GZMK, CXCR4, CCL4, CCL5, ANXA1* and several ribosomal genes compared to skin. There were also striking differences in the frequency of CD8+ T_RM_ cells between the skin and the ST: a median of 81% of skin CD8+ T-cells resided in T_RM_ cell clusters compared to 41% of ST (**Figure 2c**). Within CD8+ T_RM_ cells, the frequency of type-17 T_RM_ cells was not statistically significantly different in skin and ST. However, CD49a+ cytotoxic type-17 T_RM_ cells were significantly increased in skin compared to ST (where they were virtually absent), whereas GZMK+ cytotoxic T_RM_ cells were significantly increased in ST compared to skin (**Figure 2d**).

**Figure 2:**
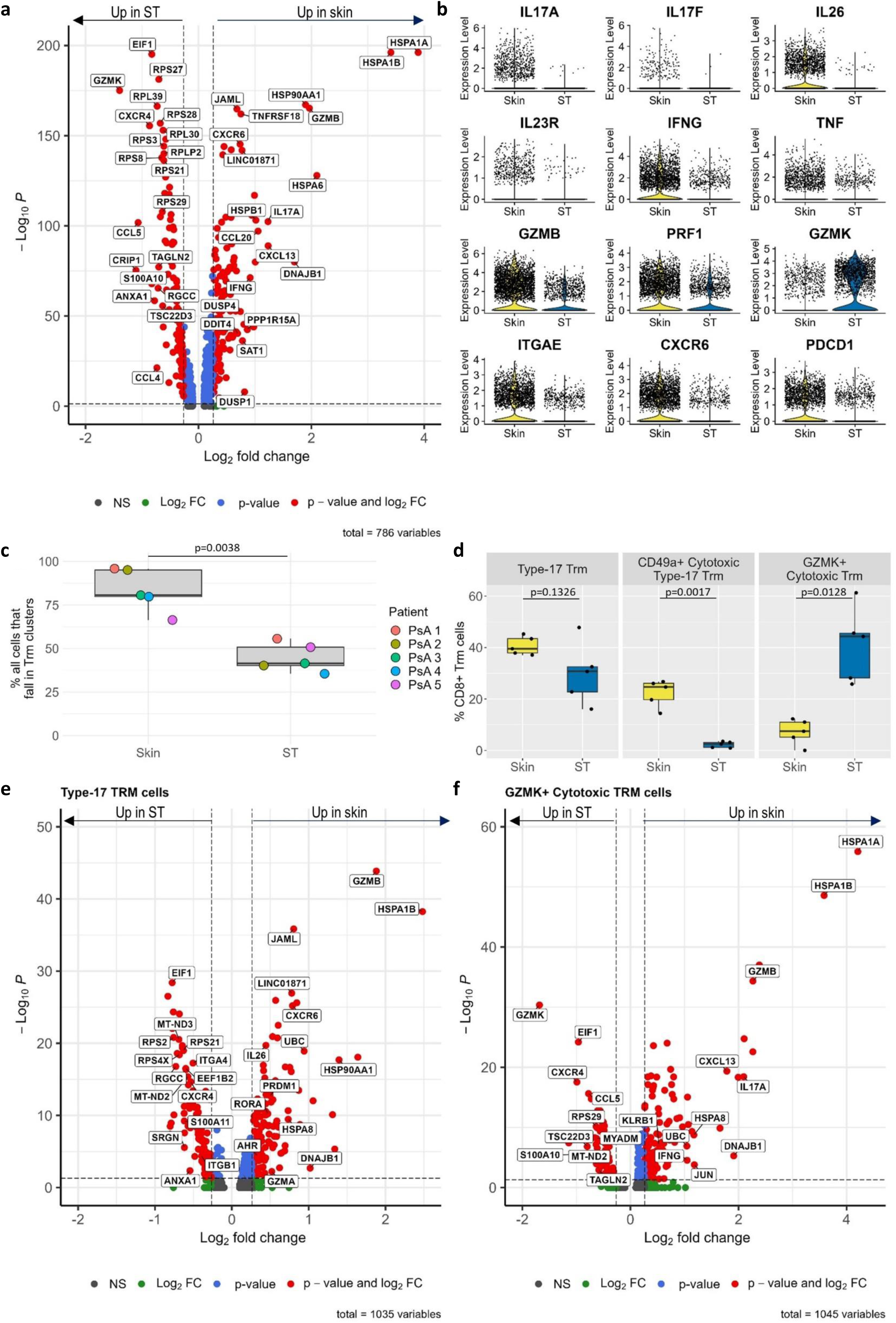
Phenotypic differences between skin and synovial CD8+ T-cells. **a)** Volcano plot depicting significant genes differentially expressed between memory CD8+ T-cells from paired samples of skin and synovial tissue from patients with PsA (n=5). Differential expression of pooled cells from all patients was calculated with the Wilcoxon signed rank test using the FindMarkers() function applied to SCTransformed RNA in Seurat. To mitigate for potential batch effects between patients or skewing of results due to patient variability, only genes which were also identified as differentially expressed by FindConservedMarkers() (which performs differential gene expression testing for each individual patient and combines the p-values using meta-analysis methods from the MetaDE R package) or were significantly differentially expressed in at least 3/5 patients after applying FindAllMarkers to each patient sample individually are labelled. The top 20 increased and decreased genes meeting these criteria are labelled. **b)** Violin plots comparing expression of specified cytokines and cytotoxic molecules in skin (n=6) and ST (n=5). **c)** % of CD8+ memory T-cells from paired skin and ST (n=5) that are located within T_RM_ cell clusters. **d)** % of CD8+ T_RM_ cells from paired skin and ST (n=5) that fall within T_RM_ clusters with each phenotype. **e,f)** Volcano plots visualising genes which are differentially expressed between paired skin and ST-cells within type-17 **(e)** and GZMK+ cytotoxic **(f)** T_RM_ cells (n=5 patients). The top 10 increased and decreased genes in addition to select genes which are specifically mentioned in the text are labelled. Differential genes calculated and criteria for labelling as described above for (**a**). Boxplots show median +/- IQR, paired t tests (two-tailed, n=5).

Even within individual CD8+ T_RM_ subsets, there were differences between skin and ST. Skin type-17 CD8+ T_RM_ cells had gene expression consistent with a stronger type-17 (*IL26*^36^*, RORA*^36^), cytotoxic (*GZMB, GZMA*) and tissue resident (*CXCR6*^34^*, PRDM1*^37^*, AHR*^38^) phenotype than ST (**Figure 2e**). In contrast, ST type-17 T_RM_ cells expressed higher levels of the chemokine receptor *CXCR4* and integrins including *ITGB1* which encodes integrin-β1 and *ITGA4* which encodes the α4 subunit of the gut homing receptor α4β7. Moreover, whilst less frequent in number, compared to those in the ST, skin CD49a+GZMK+CD8+ T_RM_ cells showed significantly increased expression of *GZMB*, potentially indicating greater cytotoxicity, and higher *IL17A,* suggesting a co-existing type-17 phenotype, whereas those from ST expressed more *GZMK* (**Figure 2f**). Skin and ST cells were not directly compared within the CD49a+ cytotoxic type-17 T_RM_ cell cluster because ST cells were virtually absent from this cluster.

These data thus show that skin CD8+ T cells overall had a stronger IL-17 signature and a higher frequency of T_RM_ cells compared to the joint. Within the T_RM_ cell clusters, the frequency of type-17 T_RM_ cells was similar in skin and ST but the majority of cytotoxic T_RM_ cells from the skin also had a type-17 signature whereas those from the joint co-expressed *GZMK* instead. Granzyme K and IL-17A were also detected at the protein level in the cell-free synovial fluid of patients with PsA suggesting that the observed gene expression translates to protein secretion in the inflamed joint (**Figure S7**).

### CD8+ T-cell clones are shared between skin and joint

Next, we sought to determine if there was evidence for clonal sharing between skin and joint. For the purpose of TCR repertoire analysis, a clone was defined as a group of T-cells expressing identical TCRα and β chains. Skin-joint shared clones were defined as T-cells from the same clone (i.e. expressing the same TCR) present in both skin and joint. We recently showed that SF and ST T-cells show considerable similarities in terms of TCR repertoire and signature (Durham et al., manuscript under review^28^), therefore SF and ST cells were pooled to increase the power to detect clones shared between the skin and the joint.

CD8+ skin-joint shared clones were detected in all six patients (**Figure 3a, b**). Across the six patients, 155 CD8+ T-cell clones were shared between the skin and the joint, comprising 1,071 CD8+ T-cells in total. In addition, some clones were shared between PsA 2, PsA 3 and PsA 6 (**Figure 3c**). HLA serotyping was not available for PsA 2 but PsA 3 and PsA 6 both were both *HLA-A*02:01*+ and therefore it is possible that clones shared between these patients may recognise antigen presented by *HLA-A*02:01* (**Table 1**). It should, however, be noted that the clones shared between these patients were different from the clones shared between the skin and joint within patients.

**Figure 3:**
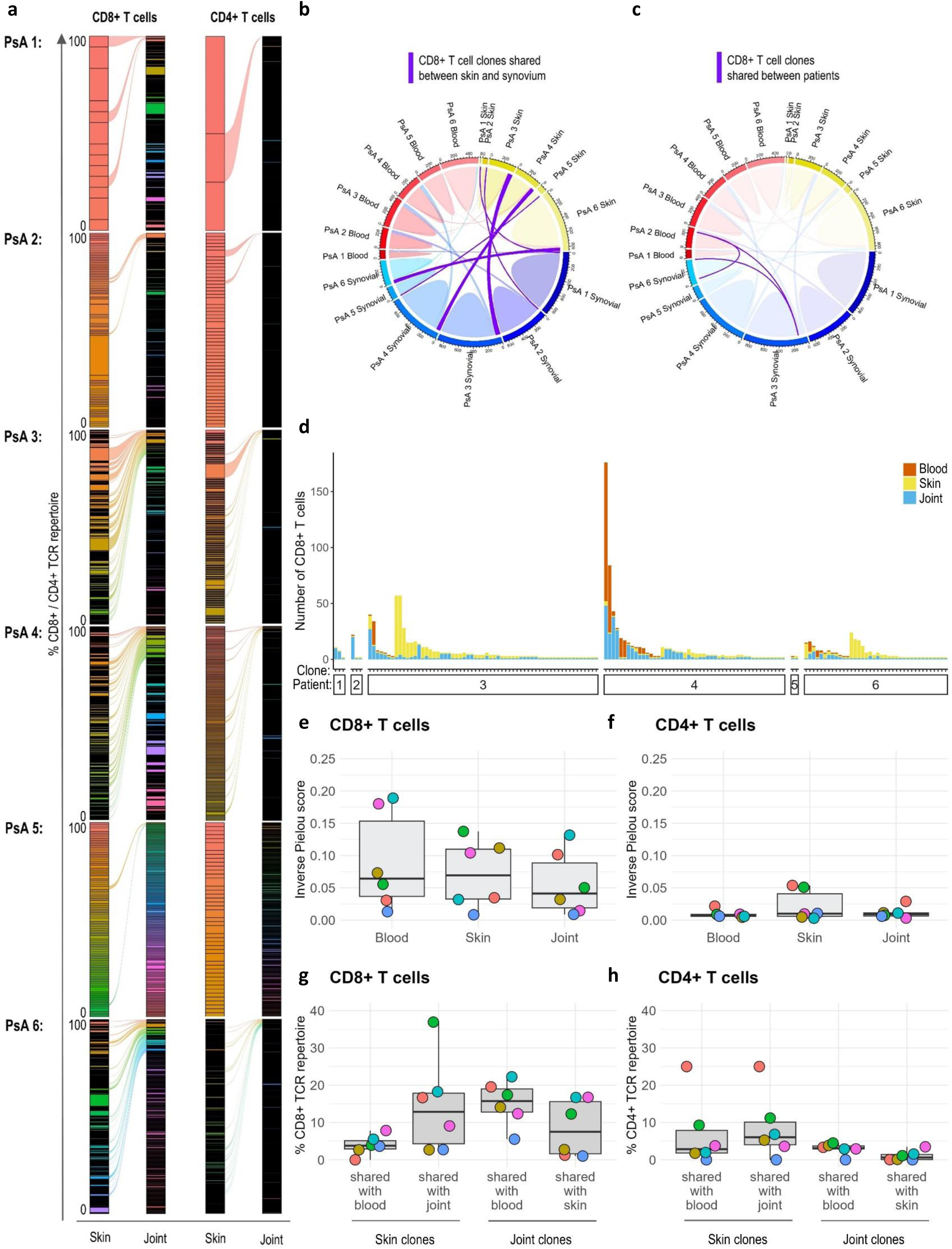
Select CD8+ and CD4+ T-cell clones are shared between skin and joint. **a)** Alluvial plots visualising entire CD8+ and CD4+ TCR repertoires in skin and joint (SF and ST combined) for each patient (n=6). Each coloured block (which is outlined in black) represents a single T-cell clone (i.e. a single/group of T-cells with the same TCR). The height of the block represents the % of the skin/joint CD8+ TCR repertoire taken up by that clone. Clones which are present in both skin and joint are represented by alluvials linking the two compartments. **b,c)** Circle plots visualising sharing of CD8+ T-cell clones between skin (yellow blocks), joint (blue blocks) and blood (red blocks) for each patient. The size of the circumferential bar for each patient is proportional to the number of single cell TCRs recovered from each tissue for that patient. **b)** Clones shared between skin and joint are represented as opaque purple lines, clones shared between skin and blood and joint and blood are represented as transparent red and blue lines respectively. **c)** Clones shared between patients (PsA 2, 3 and 6) are highlighted with purple lines. **d)** Bar chart visualising the size (number of cells contained within the clone) and tissue location of each of the 155 CD8+ T cell clones which were detected in both skin and the joint. Each vertical bar represents a CD8+ T cell clone that is shared between the skin and the joint. The height of the bar represents the number of CD8+ T-cells within that clone and the colour represents the tissue that the cells were detected in (red = blood, yellow = skin, blue = joint). Clones are split by patient and within each patient clones are ordered by size. **e,f)** Inverse Pielou scores for blood, skin and joint CD8+ **(e)** and CD4+ **(f)** T-cells. Inverse Pielou score is a measure of clonality and ranges from 0 which indicates a highly polyclonal population, to 1 which indicates a monoclonal population. **g,h)** Percentage of skin and joint CD8+ **(g)** and CD4+ **(h)** TCR repertoire which is taken up by shared clones.

Of the 155 CD8+ T cell clones that were shared between the skin and the joint, 36 clones were also detected in the blood (“triple shared clones”, **Figure 3d**). In PsA 4 (who had received a single dose of bimekizumab in the week prior to participating in the study) some of the triple shared clones were highly expanded in blood. For example, the most expanded triple shared clone comprised 124 CD8+ T cells from the blood, 48 from the joint and 4 from skin. However, most of the 36 triple shared clones contained more cells located within the skin and the joint than in the blood. Despite the presence of shared clones, the majority of clones in all patients were only detected in a single tissue and across most patients there were clones that were highly expanded in one tissue but not detected at all in others (**Figure S8**). This suggests that the detection of clones shared between tissues is not simply related to the increased probability of capturing the highly expanded clones.

There was also sharing of clones between skin and joint within the CD4+ T-cell compartment, albeit to a lesser extent (**Figure 3a**). A potential reason for reduced clone sharing by CD4+ T-cells is that CD4+ T-cells are more polyclonal than CD8+ T-cells^39^ (compare **Figure 3f** vs. **3e**). Therefore, the likelihood of capturing two cells belonging to the same clone is lower for CD4+ than for CD8+ T-cells. Overall, a median of 13% of the skin and 8% of the synovial CD8+ TCR repertoire and 6% of the skin and 1% of the synovial CD4+ TCR repertoire was made up of shared clones (**Figure 3g, h**).

We sought to characterise the signatures of shared clones by mapping them back on to the UMAP to identify which clusters they were associated with (**Figure 4a-c)**. In **Figure 4a**, the circled orange and green dots represent two different clones that are present in both skin and joint in one patient. Cells which belong to those clones are shown in orange or green on the UMAPs in **Figure 4b** and **c**, respectively. The identity of the clusters that the orange and green clones are located in can be determined by comparing their location on the UMAP to the clusters depicted in **Figure 4d**. Extending this approach to all shared clones revealed that CD8+ T-cells belonging to shared clones tended to have a similar signature in the skin and the joint, with the majority located in clusters with a T_RM_ or GZMK+ signature (**Figure 4e**). Cells from 31% of shared CD8+ clones had exactly the same signature in the skin and joint (defined as all cells of that clone falling within the same meta-cluster in both the skin and joint), 33% clones had a similar signature in the skin and the joint (defined as clones with at least one cell from the skin and one from the joint falling within the same meta-cluster) and 36% had completely different signatures (**Figure 4f**).

**Figure 4:**
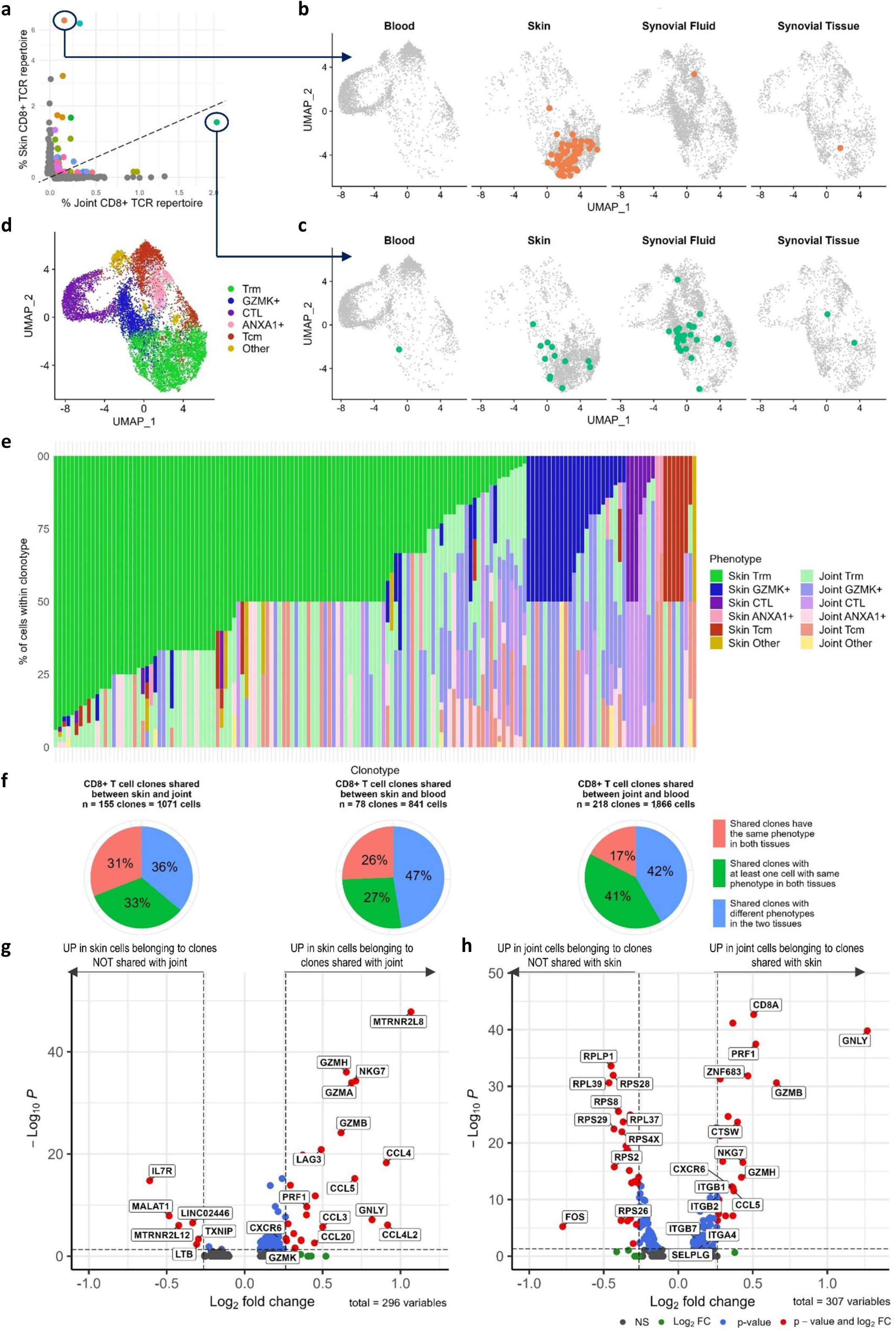
Signature of CD8+ T cell clones shared between skin and joint. **a)** Scatterplot showing frequency of CD8+ T-cell clones in skin and joint from one patient (PsA 3). Each dot represents a T-cell clone and the position of the dot along the x and y axes indicates frequency of clone in joint and skin respectively. **b,c)** Orange **(b)** and green **(c)** circled clones from **(a)** highlighted on the UMAPs to show cluster location of cells from each clonotype in each tissue. **d)** UMAP coloured by phenotype. Clusters are grouped into metaclusters with similar phenotypes as defined in Figure 1d. Clusters not defined as T_RM_, GZMK+, CTL, ANXA1+ or T_CM_ cells are grouped together as “other”. **e)** Bar chart depicting gene signature and tissue origin of cells within each of the 155 CD8+ T-cell clones shared between the skin and joint. Each vertical bar represents a CD8+ T cell clone. Colour of the bar represents the frequency of cells from each phenotype and tissue within that clone. **f)** Pie charts indicating the similarity of the signatures of clones shared between skin and joint (n=155 clones representing 1,071 CD8+ T-cells), skin and blood (n=78 clones, 841 cells) and joint and blood (n=218 clones,1,866 cells). Clones defined as having the same signature in both compartments (all cells for that clonotype are located within the same meta-cluster, in red), different signatures in the two compartments (cells for that clonotype are located within different meta-clusters in the two compartments, in blue) or similar signatures (defined as at least one cell located within the same meta-cluster from each compartment, in green). **g)** Volcano plot showing differentially expressed genes between skin cells expressing TCRs which are shared with joint (shared clones) vs. skin cells expressing TCRs which are not shared with joint (non-shared clones). **h)** Volcano plot showing differentially expressed genes between joint shared clones vs. joint non-shared clones. Low frequency of shared clones in some patients limited the utility of FindConservedMarkers() therefore, differentially expressed genes were calculated with the Wilcoxon signed rank test using FindMarkers() in Seurat which pools cells from all patients.

Shared clones in both the skin and the joint had significantly increased expression of genes associated with cytotoxicity (*GZMB, GZMH, GNLY, PRF1, NKG7)* compared to CD8+ T-cells belonging to non-shared clones (**Figure 4g, h**, **S9**). Joint shared clones also had upregulation of *CXCR6* and the transcription factor *ZNF683* (which encodes HOBIT) both of which are associated with tissue residency, suggesting a stronger T_RM_ phenotype in shared clones present in the joint compared to non-shared clones (**Figure 4h**). The integrins *ITGB1* and *ITGB2* were also significantly increased in joint cells belonging to shared clones (**Figure 4h**). We also investigated the expression of the skin homing receptor CLA (encoded by *SELPLG*) and the gut homing receptor α4β7 (encoded by *ITGA4, ITGB7*) in shared vs. non-shared clones. The expression of both receptors was significantly increased in joint shared clones compared to non-shared clones however, the fold change was low (<1.2 **Figure 4h**).

### CD4+ T_RM_ cell frequency and subset composition in the skin and the joint is different from CD8+ T_RM_ cells

In contrast to CD8+ T-cells, comparison of overall gene expression by CD4+ T-cells from lesional skin epidermis with those from inflamed ST revealed no difference in *IL17A* expression. Instead, there was significant upregulation of *FOXP3, IL2RA* and *CTLA4* in the skin compared to the joint, suggestive of a stronger T_REG_ signature (**Figure 5a, b, Supplementary Data 4**). Seurat clustering yielded 17 clusters (**Figure S10a**). Similar to CD8+ T-cells, skin and blood CD4+ T-cells clustered separately from each other on the UMAP whereas ST and SF T-cells were spread more diffusely across the UMAP indicating a broader range of signatures within CD4+ T-cells from the joint than those from either blood or skin (**Figure S10b**).

**Figure 5:**
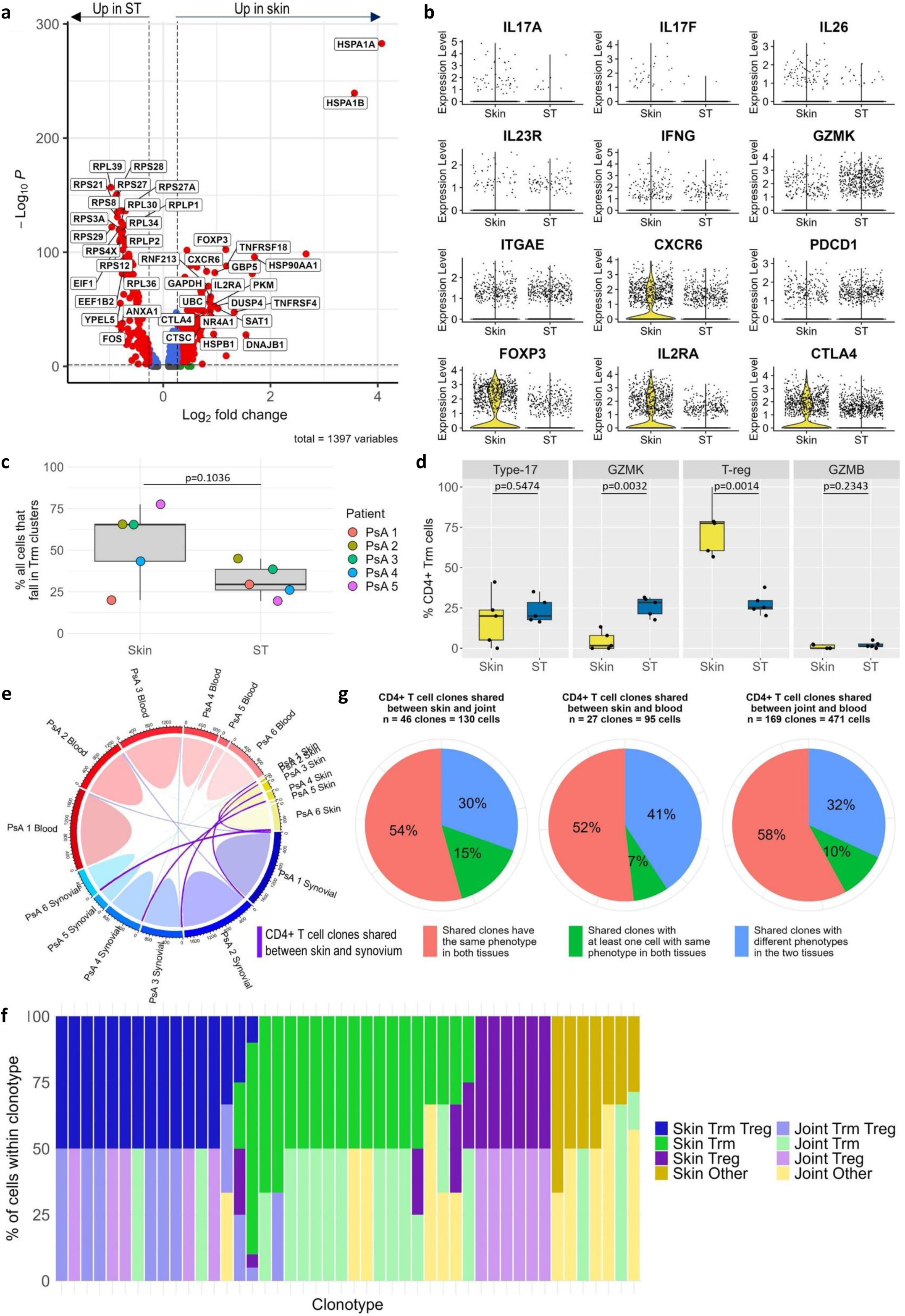
Signature of memory CD4+ T-cells in PsA: **a)** Volcano plot depicting significant genes differentially expressed between memory CD4+ T-cells from paired samples of skin and ST from patients with PsA (n=5). Differential expression of pooled cells from all patients calculated with Wilcoxon signed rank test using the FindMarkers() function applied to SCTransformed RNA in Seurat. Top 20 increased and decreased genes that were also identified as significantly differentially expressed by FindConservedMarkers() or were differentially expressed in at least 3/5 patients after applying FindAllMarkers to each patient sample individually are labelled. **b)** Violin plots comparing expression of specified genes in skin (n=6) and ST (n=5). **c)** % of CD4+ memory T-cells from paired skin and ST (n=5) located within T_RM_ cell clusters. **d)** % of CD4+ T_RM_ cells from paired skin and ST (n=5) that fall within each T_RM_ cell subset. **e)** Circle plot visualising sharing of CD4+ T-cell clones between skin epidermis (yellow blocks), joint (blue) and blood (red) for each patient. Size of circumferential bar for each patient is proportional to the number of single cell TCRs recovered from each tissue for that patient. Clones shared between skin and joint represented as opaque purple lines. Clones shared between skin and blood and joint and blood represented as transparent red and blue lines respectively. **f)** Bar chart depicting the gene signature and tissue origin of cells within each of the 46 CD4+ T-cell clones that are shared between the skin and joint. For clarity, clusters which are not defined as either T_RM_, T_RM_ T_REG_ or T_REG_ grouped as “other”. **g)** Pie charts indicating the similarity of the signatures of CD4+ T-cell clones shared between skin and joint (n=46 clones representing 130 CD4+ T-cells), skin and blood (n=27 clones, 95 cells) and joint and blood (n=169 clones, 471 cells). Clones defined as having either the same, similar or different signatures in each compartment as described in legend of Figure 4f. Boxplots depict median +/- IQR. Paired t tests (two-tailed, n=5).

Analysis of differentially expressed genes between clusters (**Figure S11, Supplementary Data 5**) enabled grouping of clusters with similar gene signatures including T_RM_ cells (6, 9, 10 and 15) and T_REG_s, both with (clusters 11 and 13) and without (clusters 2 and 12) a T_RM_-like signature (**Figure S10c, d**). CD4+ T_RM_ cell clusters were identified and characterised using the same approach as for CD8+ T-cells (**Figure S10e-g**). Unlike CD8+ T-cells, there was no significant difference in the frequency of CD4+ T_RM_ cells in the skin and ST (**Figure 5c**). Examination of gene expression (**Figure S10f**) and GSEA comparing each CD4+ T_RM_ cluster to pooled cells from other T_RM_ clusters (**Figure S10g**) identified four T_RM_ subsets: type-17 (cluster 9), CD49a+*GZMK*+ (cluster 6), T_REG_s (clusters 11, 13) and a small cluster of CD49a+*GZMB+* T_RM_ cells (cluster 15). There was also a T_RM_ cluster with a mixed signature (cluster 10). The frequency of type-17 CD4+ T_RM_ cells was similar in the skin and ST whereas CD49a+GZMK+ T_RM_ cells were increased in ST, and T_REG_ T_RM_-like cells were increased in skin (**Figure 5d**).

Similar to CD8+ shared clones, we detected shared CD4+ T-cell clones between the skin and joint, albeit lower in frequency: 46 clones were shared comprising 130 cells in total, (**Figure 3a**, **Figure 5e**). Most shared clones had a T_RM_ or T_REG_ T_RM_-like signature and as with CD8+ T-cells, cells belonging to CD4+ shared clones tended to have the same signature in the skin and the joint (**Figure 5f, g**).

## Discussion

By comparing paired skin and joint samples from individuals with psoriatic arthritis we demonstrate shared T-cell clonality and signatures between these sites. We identified a stronger T_RM_ and IL-17 signature within CD8+ T-cells from the skin compared to the joint and higher expression of *GZMK* in CD8+ T-cells from the joint. Despite differences in the overall signature of CD8+ T-cells from the skin and joint, the shared clones tend to have similar signatures in both sites. Shared CD8+ T-cell clones in both the skin and joint had significantly increased expression of genes associated with cytotoxicity (*GZMB, GNLY, PRF1, GZMH, NKG7)* compared to non-shared clones. This could indicate increased pathogenic potential of these shared clones, and suggests that they are more activated than their non-shared counterparts and therefore more likely to drive inflammation in both sites.These data indicate that the CD8+ T-cell infiltrate in skin and joint is linked in terms of T-cell clonality and raise the possibility that these cells migrate between the skin and the joint to propagate inflammation across both sites.

The huge diversity of the human TCR repertoire is a result of somatic recombination of V(D)J genes and introduction of random nucleotides at junctions between gene segments during T-cell development^40^. It is highly unlikely that two independent T-cell precursors would rearrange their TCR into the exact same sequence. Thus, two T-cells expressing the same TCR sequence in skin and joint in PsA will almost certainly be descended from the same parent T-cell. Most clones that were shared between the skin and joint were not detected in blood, and those that were, were usually present at higher frequencies in the skin/joint than in the blood. This suggests that the presence of skin-joint shared clones is not simply because clones which were highly expanded in blood seeded in multiple tissues by chance. Our finding that skin-joint shared clones exist could indicate that the same antigen is present at both sites, and/or that T-cells migrate between the skin and the joint. Indeed, whilst T_RM_ cells are typically described as permanently resident within tissue, a subset can egress from tissue and travel in the blood to establish residency at distant sites or even trans-differentiate into different memory subtypes^41–46^. For example, in humans, “ex-skin T_RM_ cells” that express CD103 and CLA can be detected in blood^44^. This may explain how inflammation spreads from site to site in multi-system immunological diseases such as PsA and provides a potential explanation for the skin-joint shared clones identified in our study.

To investigate whether the clones that were shared between the skin and the joint exhibited features associated with increased potential to for tissue homing, we compared the expression of adhesion and tissue homing molecules by shared vs. non shared clones. Compared to non-shared clones, shared CD8+ T-cell clones in the joint expressed significantly higher levels of *ITGB1 and ITGB2* (which encode integrin-β1 and integrin-β2 respectively). Integrin-β1 and integrin-β2 form heterodimers with multiple different α-integrins to function as adhesion receptors. They govern the cells’ interaction with the extracellular matrix and with other cells, providing mechanical support and modulating T-cell motility, activation, proliferation and differentiation^47^. Integrin-β1 expression is also associated with increased cytotoxic capacity of CD8+ T-cells^48^. The genes encoding the skin homing receptor CLA (*SELPLG* which encodes PSGL-1 which is post-translationally modified to CLA) and the gut homing receptor α4β7 (*ITGA4* and *ITGB7*) were also marginally increased in CD8+ T cell clones in the joint that were shared with skin compared to non-shared clones. However, the fold change of these genes was low and therefore further work, including the investigation of their expression at the protein level in shared vs non-shared clones is required. Whether the increased expression of integrins (and/or skin/gut homing receptors) by CD8+ shared clones in the synovium is evidence of their enhanced motility and capacity to migrate between sites, or is reflective of their increased activation state and pathogenicity, is currently not known. However, the presence of activated, cytotoxic T_RM_ cells in the joint which share TCR sequences with their counterparts in the skin indicates that the T-cell infiltrate in the skin and synovium is linked in terms of T-cell clonality, and supports the concept that T-cells migrate between the skin and the joint.

Two putative autoantigens have been reported in psoriasis (the melanocyte peptide ADAMTSL5^49^, and antimicrobial peptide LL37^50^), both of which are presented by the psoriasis-associated HLA-C*06:02, activate T-cells in psoriatic skin, and can induce type-17 responses. Moreover, the disease-associated haplotype of ERAP-1 enhances presentation of ADAMTSL5 leading to enhanced activation of ADAMTSL5-specific CD8+ T-cells^51^. A recent study of patients with Ankylosing Spondylitis (AS), which shares many clinical, genetic and immunological characteristics with PsA, has identified shared T-cell clones, most of which were CD8+, between the eye and joint, two sites of disease in AS^52^. Detailed analysis of these clonal TCR and HLA-B27 interactions, identified potential self and/or microbial antigens that might drive both joint and eye inflammation in AS^52^. These data create a strong rationale to investigate (self-)antigen driven recruitment and/or activation of T-cells in the skin and joint in PsA. However, the migration of T-cells between the skin and the joint may not require the same antigen to be present in both sites: studies in mice have shown that antigen-independent inflammation in tissue is sufficient to drive T-cell recruitment and T_RM_ cell differentiation^44,53^. Thus, inflammation of a joint arising from biomechanical injury may non-specifically recruit autoreactive ex-skin T_RM_ T-cells from the blood into the joint where they could take up residence and trigger the development and persistence of PsA.

Despite the enrichment of IL-17+CD8+ T_RM_ cells in both skin and joint, T-cells belonging to skin-joint shared clones did not have increased expression of type-17 associated genes compared to those belonging to non-shared clones. This is surprising since in mice, type-17 T_RM_ cells exhibit a greater propensity to egress from skin than cytotoxic T_RM_ cells^54^. Our finding could be reflective of inherent differences between this particular mouse model and human psoriatic tissue, or of the high degree of plasticity inherent to type-17 cells. For example, studies in mice have shown that upon migration to sites of inflammation, IL-17+CD8+ T-cells upregulate expression of IFNγ and lose/reduce expression of IL-17 to become type-1 cells^55–59^.

The finding that skin CD8+ T-cells have a stronger IL-17 signature than those from the joint correlates with clinical trial data and real-world experience that IL-17 inhibition produces a more profound improvement in psoriatic skin inflammation than in joint inflammation^9,10^. It is consistent with previous studies comparing skin and synovium by bulk RNA sequencing which reported stronger IL-17/IL-23 signatures in the skin^25,26^. Despite the stronger IL-17 signature in the skin, CD8+ type-17 T_RM_ cells are present in the joint at frequencies consistent with previous work^6,7^. Our findings therefore suggest that whilst IL-17 may be a *dominant* driver of inflammation in the skin, other factors, *in addition to* IL-17, contribute to driving inflammation in the joints. The apparent redundancy in the pathways driving synovial inflammation in PsA may explain why blocking IL-17 alone has a less dramatic effect at reducing joint inflammation than skin inflammation. Treatment of refractory PsA may require combination therapy with drugs that block multiple cytokines (for example, a monoclonal antibody that blocks IL-17 signalling in combination with anti-TNF) and/or new targets.

One new target may be GZMK. The presence of CD8+GZMK+ T-cells has been reported in inflamed synovium in RA, inflamed bowel in Crohn’s disease and ulcerative colitis and in labial glands in primary Sjogren’s syndrome^60,61^. Thus, CD8+ GZMK+ T-cells may have a role in driving inflammation in multiple diseases and across multiple tissues. Unlike granzyme B, granzyme K does not cleave intracellular caspases to induce apoptosis and does not require perforin for its action^60^. Instead, it cleaves extracellular proteins on fibroblasts to trigger activation of intracellular signalling pathways leading to production of inflammatory mediators including IL-6 and CCL2^60^. Our data therefore supports investigation into the viability of blocking granzyme K to treat synovial inflammation in PsA.

Our analysis also identified four subsets of CD4+ T_RM_ cells: 1) CD4+ Type-17 T_RM_ cells, which are present at equal frequencies in the skin and the joint, 2) CD4+ CD49a+GZMK+ T_RM_ cells, which are increased in the joint, 3) CD4+ T_REG_ T_RM_-like cells, which are enriched in the skin, and 4) CD4+ CD49a+GZMK+ T_RM_ cells which are present at very low frequencies in both sites. This is an extension of our previous work using CyTOF in which we identified just one cluster of CD4+ T_RM_ cells based on co-expression of CD103 and CD69^7^. The likely reason is that single cell RNA sequencing employed in the current study enabled the use of GSEA to identify additional CD103-CD4+ T_RM_ cells. Further characterisation of these cells to identify surface markers will allow better understanding of their role in human disease. In humans, CD4+ T_RM_ cells tend to have a higher propensity to egress from skin and migrate to distant sites than CD8+ T_RM_ cells^44,46^. Studies in mice have shown that tissue T_REG_ cells readily recirculate between the vasculature and the skin^62^. Furthermore, a recent study of patients with JIA showed that shared CD4+ T_REG_ clones can be detected in different joints of the same patient, suggesting that they can migrate between tissues in humans^63^. The majority of shared CD4+ T cell clones in our dataset had a T_RM_ or T_REG_ signature which supports the hypothesis that CD4+ T_RM_ and/or T_REG_ cells can also migrate between the skin and the joint. In this study we investigated the inflammatory T-cell infiltrate in skin epidermis. Further work comparing the TCR repertoire of full thickness skin (since the majority of CD4+ T-cells are located in the dermis) with the joint may lead to the identification of greater numbers of shared CD4+ T-cell clones to investigate this in more depth.

Our study has some limitations. We enrolled six patients for this study to donate paired biopsies from inflamed skin and an inflamed joint for a research indication and in a non-interventional context. The patients had different duration of disease and were on different treatments. Whilst every effort was made to mitigate for patient variation and/or batch effect, increased patient numbers will be required to confirm that the results can be generalised to the population of patients with PsA. Furthermore, whilst all 5 patients for whom HLA serotyping was available had PsA and/or psoriasis associated alleles, we did not recruit patients with a specific HLA haplotype. In this study we performed single cell RNA sequencing on memory T-cells dissociated from tissue. Tissue dissociation can introduce artefact^28,35^ and removes spatial information. Spatial transcriptomics can mitigate both limitations, however, at the time of this work, platforms for spatial transcriptomics did not support TCR sequencing. Therefore, the current approach was the most appropriate option to investigate the presence of shared T-cell clones between skin and synovium.

In summary, we show that T-cells expressing identical TCRs are present in the skin and the joint of patients with PsA and that CD8+ shared clones tend to have common signatures and increased pro-inflammatory potential at both sites. This could indicate that T-cells are migrating between the skin and the joint. This potential migration might be driven by a common antigen at both sites, in which case antigen-specific immunotherapy may have a role for inducing long-term tolerance to the antigen(s). Alternatively, T-cell migration might be antigen-independent, in which case blocking chemokine signalling may prevent T-cell migration. Further sophisticated investigation will be required to explicitly determine whether T-cells migrate between the skin and the joint in humans and the mechanisms by which this is happening. Nonetheless, the demonstration that skin and joint inflammation in PsA is linked through the presence of phenotypically similar T-cell clones, will inform novel translational approaches towards treatment of PsA.

## Methods

### Participant recruitment

Participants fulfilling the Classification of Psoriatic Arthritis (CASPAR) study group criteria^64^ were recruited from Guy’s and St Thomas’ NHS Foundation Trust, London, UK. **Table 1** gives demographic and clinical information for each patient. Participants donated synovial tissue (ST) and/or synovial fluid (SF) from an inflamed knee, one 6mm skin punch biopsy from lesional psoriasis and blood. Five patients donated ST by ultrasound guided needle biopsy during which 6-9 biopsies, each measuring ∼1x1x2mm, were taken for this study. One patient (PsA 1) donated ST at the time of knee replacement surgery.

### Cell isolation

In the five patients where ST was obtained by ultrasound-guided needle biopsy, samples were processed immediately and cells were loaded onto the 10x Genomics chip within 8 hours of collection. Peripheral blood and SF mononuclear cells (PBMC and SFMC respectively) were isolated by density gradient centrifugation using Lymphoprep^TM^. 6-9 ST biopsies were placed directly into serum-free RPMI media containing 0.3mg/ml Liberase and 0.1mg/ml DNase and processed using a custom 30-minute programme on the gentleMACS dissociator (Miltenyi Biotech) which combines enzymatic and mechanical digestion to extract cells from tissue. The reaction was then quenched by the addition of 2.5ml RPMI media supplemented with 10% fetal bovine serum and 1% penicillin, streptomycin and L-glutamine and samples were strained. Skin punch biopsies were incubated at 37°c in 10mg/ml Dispase II in Hanks Balanced Salt Solution for one hour. The epidermis was then peeled off and processed using the gentleMACS dissociator as described above for ST.

In the patient where ST was obtained at knee replacement surgery, ST was minced into ∼1-2mm^3^ sized pieces and PBMC and SFMC were isolated and skin epidermis separated from the dermis as described above. Samples were then cryopreserved in culture media containing 90% FCS and 10% DMSO in liquid nitrogen until use. Upon thawing, all subsequent steps were the same as those described above.

### Sample staining

Following cell isolation, PBMC, SFMC and ST and skin epidermis digests underwent identical protocols for staining and sorting. Cells were transferred to LoBind Eppendorfs, washed in PBS, resuspended in 50ul/staining volume of Fc block mastermix (5ul Fc block and 45ul cell staining buffer, both Biolegend; one staining volume = up to 2 x 10^6^ cells) and incubated at room temperature for 10 minutes. Following this, 49ul/staining volume of staining mastermix (containing fluorescently labeled antibodies and Cellular Indexing of Transcriptomes and Epitopes by sequencing (CITE-Seq) antibodies, **Supplementary Table 1**) and 1ul/staining volume of hashtag TotalSeqC antibody mastermix (Biolegend, **Supplementary Table 2**) was added to each sample and cells were incubated for 30 minutes at 4°C. For each patient, the two samples with the highest cell number (for PsA 1, 2, 3 and 6 this was PBMC and SFMC and for PsA 4 and 5 this was PBMC and ST) were stained in duplicate to enable the use of 6 separate hashtags across the 4 samples to enhance doublet detection. Cells were then washed twice in cell staining buffer and a third time in PBS containing 1% FCS, then resuspended in 300ul PBS containing 1% FCS and transferred to sterile FACS tubes. Replicate cells from the same tissue which were stained with different hashtags were pooled at this point.

### Fluorescent-Activated Cell Sorting

Memory T-cells (CD45RA-CD27+, CD45RA-CD27+ and CD45RA+CD27-) were sorted on BD Aria cell sorter using an 85um nozzle for ST and skin epidermis digests and a 70um nozzle for PBMC and SFMC. Gating strategy is shown in **Supplementary Figure 1**. DAPI was added to label non-viable cells immediately prior to sorting. Cells from all four tissue compartments were pooled in a single LoBind Eppendorf that had been pre-coated with FCS and were retained on ice pending library preparation.

### Single cell library preparation and sequencing

A maximum of 20,000 sorted memory T-cells per well were immediately loaded onto the 10x Genomics chip. Single cell libraries were created using the Chromium Single-Cell 5’ Reagent Kits v1.1, Libraries were sequenced on the NextSeq 2000 platform.

### Pre-processing of single cell RNAseq data and QC

Raw reads were aligned to the human transcriptome and “multi” option of CellRanger software package (v6.1.1) with default parameters. Outputs from CellRanger were loaded into Seurat (version 4.3.0). TCR genes were removed from the RNA assay and added to the Seurat object as a separate assay (“TCR assay”). This was to prevent TCR genes having any effect on sample integration or cell clustering. Samples were demultiplexed (using MULTIseqDemux()). Poor quality cells (defined as cells with very high or low numbers of genes (nFeature_RNA), very high or low number of absolute RNA counts (nCount_RNA), a high percentage of mitochondrial genes or absolute ADT counts > 10,000 (which likely indicates the presence of a clump of CITE-seq antibodies)) were removed. The thresholds for nFeature_RNA, nCount_RNA and %mitochondrial QC metrics were defined as three times the mean absolute deviation of each metric in each independent sample^65^. The advantage of calculating automatic thresholds for each sample rather than using a “one size fits all” threshold for all samples is that automatically calculated thresholds take into account variation between samples. Doublets, which were identified by MULTIseqDemux and/or cells which expressed more than one TCRβ gene or more than two TCRα genes, were also removed. An additional QC step was performed for PsA 5 which, despite high cell viability during cell sorting, had higher ambient RNA in the gene expression library. For this sample, only cells which had paired TCR sequences (and therefore most likely represented true, viable cells) were taken forward to QC.

### Single cell RNAseq analysis

Single cell RNAseq data was analysed in R (version 2.2.2) using Seurat (version 4.3.0). The 7 datasets (two libraries were created for PsA 3, one library for each of the other 5 patients) were individually normalised using SCTransform v2 with regression of cell cycle score, the percentage of mitochondrial genes and the digestion module score^28^ (RNA assay) and NormalizeData() and ScaleData (ADT and TCR assays) and integrated^29,31^. The digestion module score was created using the list of 512 genes which comprised the “core digestion signature” reported in ^35^. Dimensionality reduction was performed using the integrated RNA assay to calculate 30 principal components and construct a SNN graph using 30 dimensions. Clustering was performed on the integrated assay using resolution 0.9 with otherwise default parameters yielding 24 clusters.

### Identification of CD8+ and CD4+ T-cells

CD8+ and CD4+ T-cells were extracted from the integrated dataset and analysed separately. Clusters were broadly classified as “CD4+” (clusters 0, 1, 3, 7, 8, 12, 13, 15 and 22), “CD8+” (clusters 2, 4, 5, 6, 10, 14, 17, 18, 19, 20 and 21) or “mixed” (clusters 9, 11, 16, and 23) based on expression of *CD4* and *CD8A* (**Supplementary Figure 3c**). However, some clusters, for example cluster 6, predominantly comprised *CD8A*+ cells but did contain some cells that expressed *CD4* (**Figure S3d**). Similarly, there were *CD4*+ clusters which contained some cells that expressed *CD8A.* Therefore, *CD8A+CD4-* cells which were located in CD4+/mixed clusters were extracted and added to the CD8+ T cell dataset. And, *CD4+CD8A-* cells which were located in CD8+/mixed clusters were extracted and added to the CD4+ T cell dataset. The separate CD8+ and CD4+ datasets were then split up into their constituent sample libraries and re-normalised, integrated and clustered (with resolution 1 and 0.9 for the CD8+ and CD4+ dataset respectively) using the same approach as described above.

### Differential Gene Expression

Wilcoxon rank sum tests were performed using the SCTransformed assays to identify differentially expressed genes for each cluster/tissue using the FindConservedMarkers(), FindMarkers() and FindAllMarkers() functions. Combined p values (FindConservedMarkers) and adjusted p-values (FindMarkers and FindAllMarkers) <0.05 were considered statistically significant. FindMarkers/FindAllMarkers pools cells from all samples in an integrated object and therefore, the results may be subject to batch effect between samples or be skewed by one sample. In contrast, FindConservedMarkers() performs differential gene expression testing for each sample (in this case, patient) separately and combines the p-values using meta-analysis methods from the MetaDE R package. For this reason, where feasible, we report differentially expressed genes that are identified by either FindConservedMarkers() or which are significant in the majority of patients when FindMarkers/FindAllMarkers is performed on each patient individually.

### Gene Set Enrichment Analysis (GSEA)

GSEA was performed using GSEA v4.2.3^66^. Lung CD4+ and CD8+ core T_RM_ signatures were obtained from Kumar et al ^34^. With regards to other gene signatures used for GSEA, signatures from psoriatic skin T-cell subsets comprised T_RM_, Tc1, Tc17 and Tc17/Tc22 signatures from ^8^, CD49a+ and CD49a- T_RM_ signatures from ^4^, and psoriatic and healthy skin type-17 CD4+ and CD8+ T_RM_ cells and psoriatic and healthy skin CD4+ and CD8+ T_REG_s from ^67^. Signatures from PsA synovial fluid T-cell subsets comprised type-17 and CD49a+GZMK+ T_RM_ signatures from our previous work^7^, CD8+ HLA-DR high T-cells from ^16^, and, canonical T_REG_s and CCR4+Helios+ T_REG_s signatures from ^32^.

### Single cell TCR sequencing analysis

TCR repertoire analysis was performed in R using the scRepertoire and circulize packages^68,69^.

### Statistical analysis

Statistical analysis was performed in R (v4.2.2) and GraphPad Prism (v10). Statistical tests, p values and number of samples are detailed in relevant figure legends. Detailed results of statistical analysis are reported in **Supplementary Data 6**.

## Acknowledgements

The authors are grateful to the patients who participated in this study and to Miss Diane Back and Mr Peter Earnshaw, Trauma and Orthopaedics department at Guy’s & St. Thomas’ NHS Foundation Trust, who assisted with ST collection from surgery. The authors acknowledge the following funding: MRC Clinical Research Training Fellowship (ref MR/P018904/1 to LED), Versus Arthritis programme grant (ref 21139 to LST), the Foundation for Rheumatology Research (FOREUM to LST), the King’s Health Partners Centre for Translational Medicine (to FH, BWK and LST) and the NIHR-BRC at KCL/GSTT.

## Author contributions

L.E.D., B.W.K. and L.S.T. conceived the project, designed the experiments, interpreted the results and wrote the manuscript. L.E.D., F.H., N.N. and B.W.K. obtained clinical samples, L.E.D., S.E.R., E.H.G., R.R., R.N., K.F. and P.D. contributed to data acquisition, L.E.D., A.M., P.D., E.H.G. and G.A.M.P. contributed to data analysis. B.W.K and L.S.T. supervised the project. All authors contributed to the writing and revising of the manuscript.

## Author disclosures

BWK has received speaker or consultancy fees and/or research support from AbbVie, Eli Lilly, Galapagos, Janssen, Novartis, Pfizer and UCB, outside of this work. LST has received speaker or consultancy fees and/or research support from AbbVie, GSK, IMID forum, Sanofi and UCB, outside of this work.

## Notes

### Summary of Updates

Table 1 has been updated, no other changes have been made to manuscript.

