## Supplementary Figures and Tables for "Clonal sharing of CD8+ T-cells links skin and joint inflammation in psoriatic arthritis"

### Fig. S1 Durham et al.

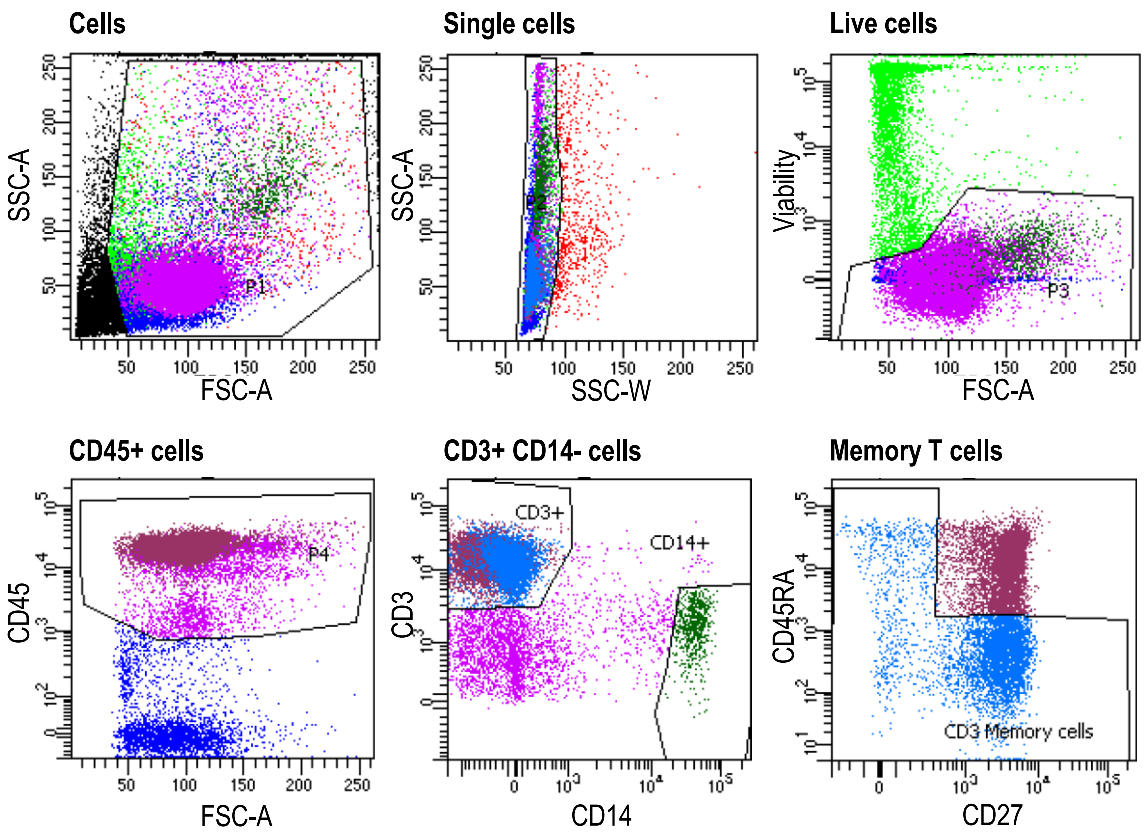


**Figure S1: Gating strategy**

Gating strategy used to sort live memory (CD45RA-CD27-, CD45RA-CD27+ and CD45RA+CD27-) T-cells from PBMC, SFMC, skin epidermis and synovial tissue digests. Cells were stained with fluorescent-conjugated antibodies and CITE-seq antibodies prior to sorting. DAPI was added immediately prior to sorting to enable exclusion of dead cells. The gating strategy is shown for cells (FSC-A vs SSC-A), singlets (SSC-W vs SSC-A), live cells (DAPI-), CD45+ cells, CD3+CD14- cells and CD45RA-CD27-/CD45RA-CD27+/CD45RA+CD27- cells. Representative example using PBMC from patient PsA 2.

### Fig. S2 Durham et al.

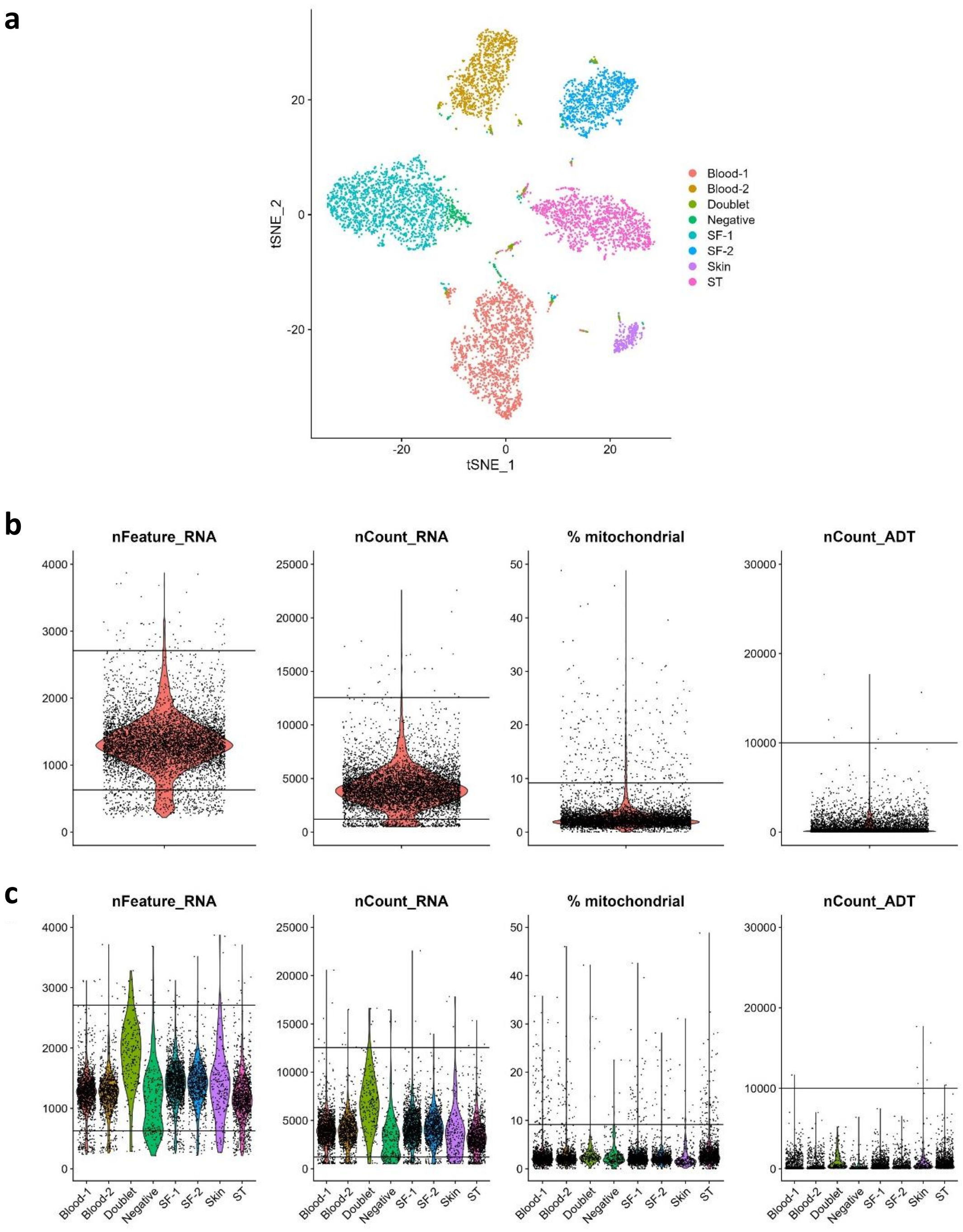


**Figure S2: Quality control of single cell sequencing**

QC plots from one representative patient (PsA 2). Samples were demultiplexed then filtered to remove poor quality cells (defined as cells with very high or low numbers of genes, very high or low number of absolute counts or a high percentage of mitochondrial genes). The thresholds for nFeature_RNA, nCount_RNA and %mitochondrial QC metrics were defined as three times the mean absolute deviation of each metric in each independent sample[65] **a)** tSNE plot showing the results of de-multiplexing the hashtags. Cells coloured by assigned identity. Blood and synovial fluid were split in half and each half was stained with a different hashtag. This was to increase the overall number of hashtags and thereby improve doublet detection. **b)** Violin plots depicting the number of genes (nFeature_RNA) and number of RNA counts detected per cell (nCount_RNA), the percent of detected genes which are mitochondrial genes (% mitochondrial) and number of ADT counts detected per cell (nCount_ADT). Horizontal lines indicate the QC thresholds for this patient. **c)** Violin plots visualising the QC metrics grouped by the identity assigned by de-multiplexing the hashtags.

### Fig. S3 Durham et al.

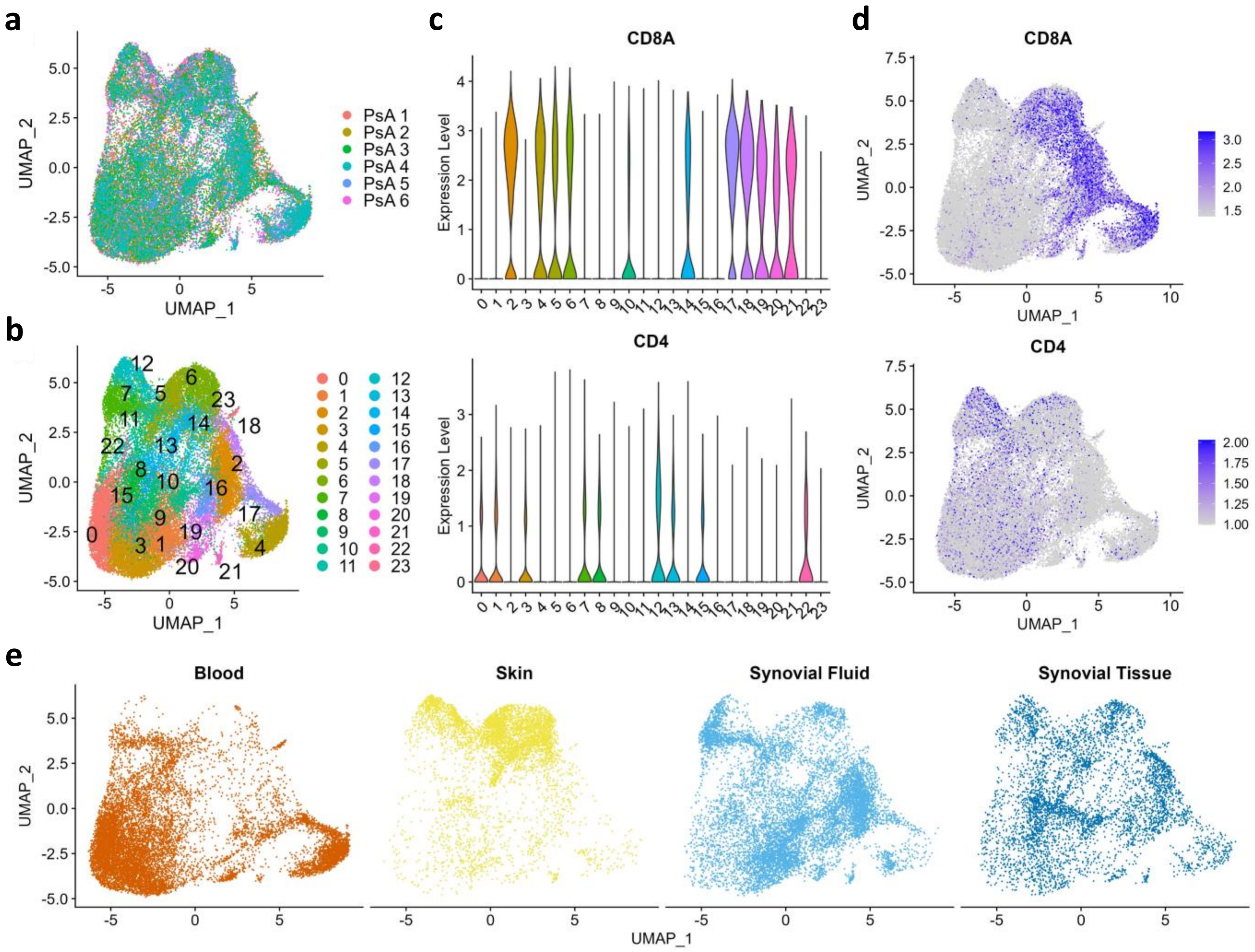


**Figure S3: Integrated analysis of 35,491 memory T cells from paired samples of blood, skin epidermis from inflamed psoriatic skin and synovial tissue and/or synovial fluid from inflamed knees from 6 patients with PsA.**

**a, b)** UMAPs with cells coloured according to **(a)** patient and **(b)** the 24 cell populations obtained after Seurat clustering. **c)** Violin plots visualising CD8A and CD4 expression within each cluster. **d)** UMAPs with cells coloured according to *CD8A* and *CD4* expression. **e)** UMAPs split by tissue of origin.

**Fig. S4 Durham et al.**

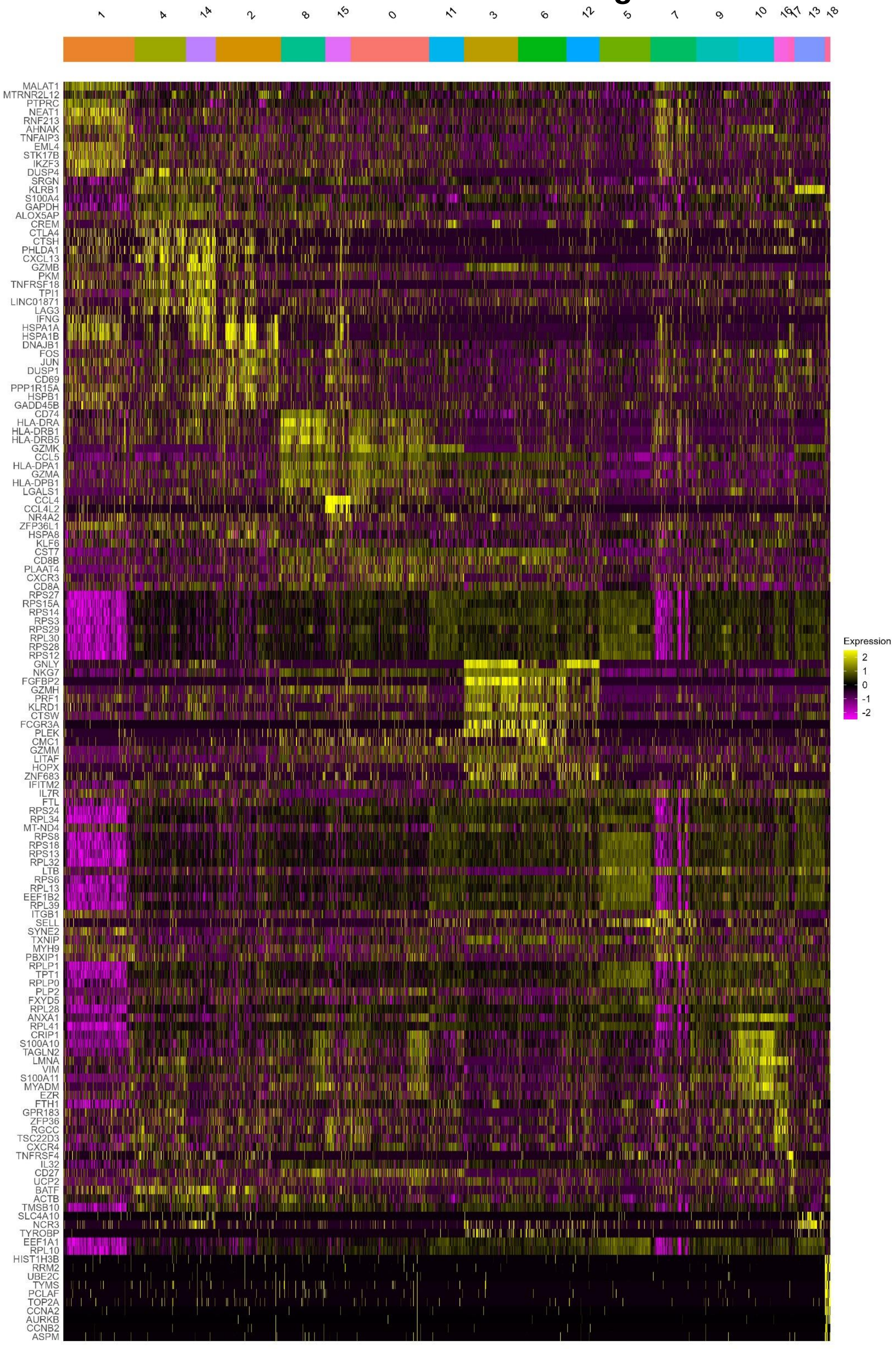

**Figure S4: Heatmap visualising the top 10 genes differentially expressed by each of the 19 CD8+ T cell clusters in the CD8+ T cell only analysis.** Clusters grouped into groups with similar phenotypes. Differential expression by SCTransformed RNA was calculated using the Wilcoxon signed rank test using the FindConservedMarkers() function Seurat. FindConservedMarkers() mitigates for potential batch effect between patients by performing differential gene expression testing for each patient separately and combining the p-values using meta-analysis methods from the MetaDE R package. Combined p value $< 0.05$ was used to identify significantly differentially expressed genes. Log2FC for each of the 6 patients was averaged and genes were ranked in order of average Log2FC change.

### Fig. S5 Durham et al.

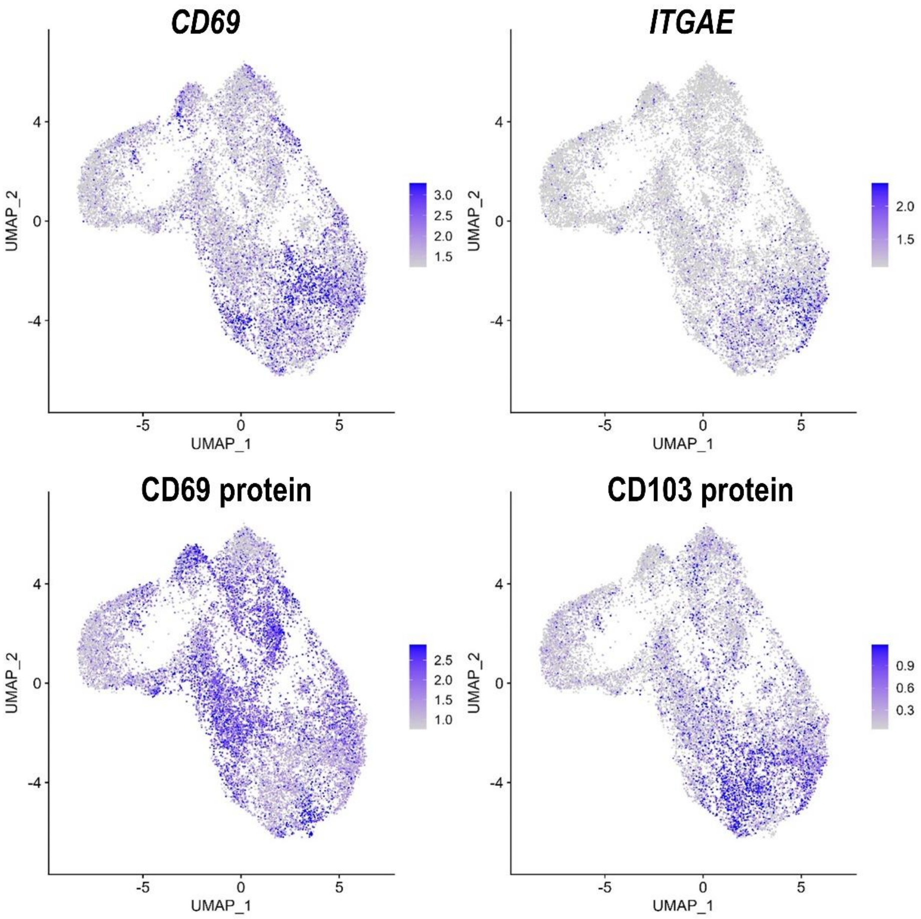


**Figure S5: Expression of CD69 and CD103 by CD8+ T cells from skin, ST, SF and blood.**
UMAPs coloured by expression of *CD69* and *ITGAE* (encodes CD103) RNA (top row) and CD69 and CD103 protein (bottom row) (n=6 patients)

**Fig. S6 Durham et al.**

a

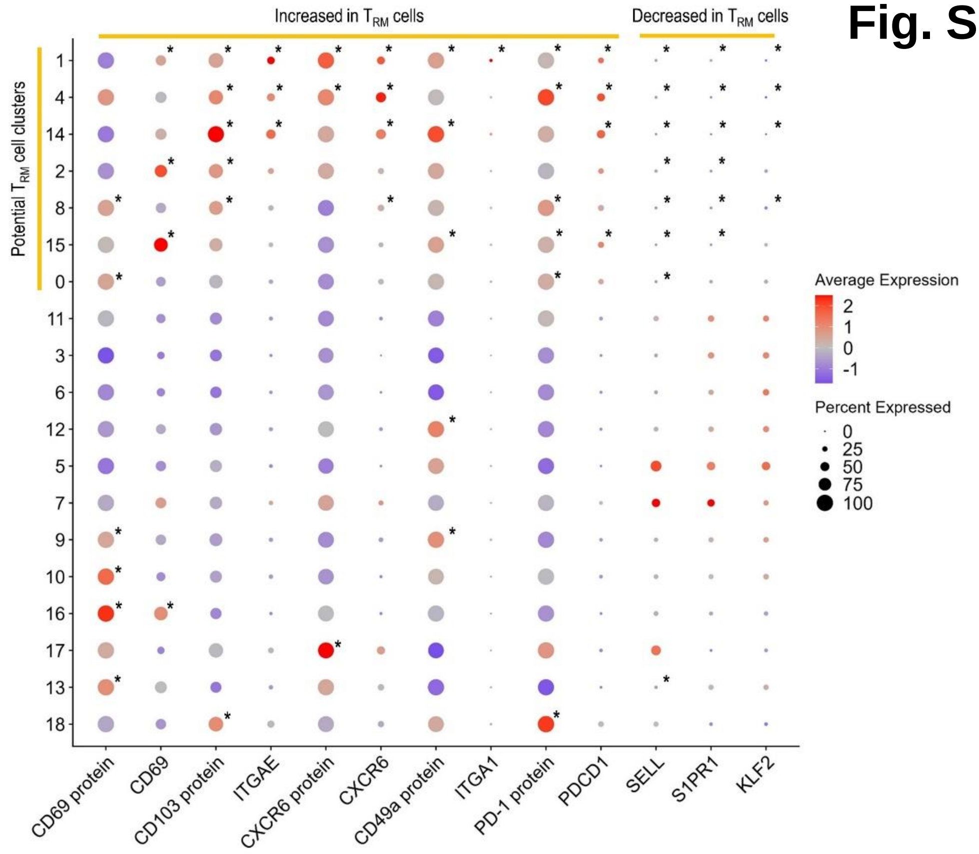


b

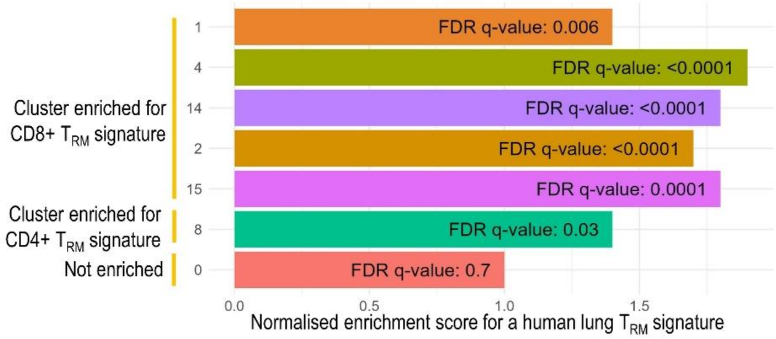


c

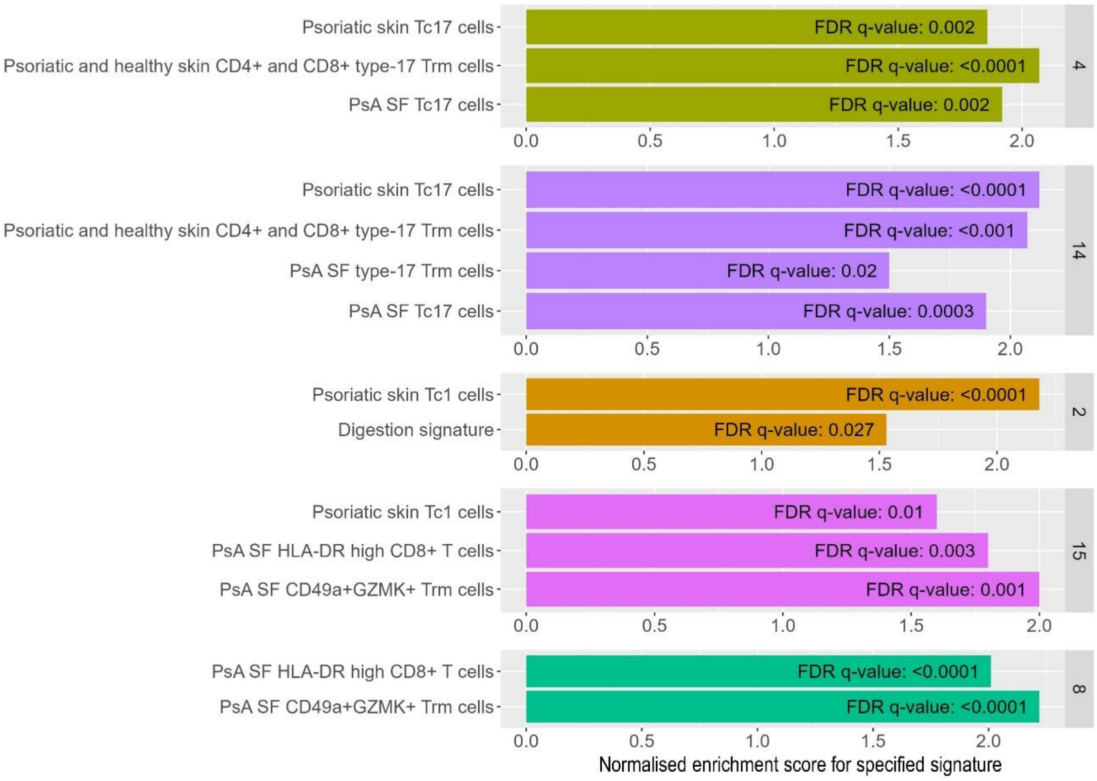

**Figure S6: Identification and characterisation of CD8+ $T_{RM}$ cell clusters**

**a)** Dot plot depicting the expression of specified genes/proteins across the CD8+ T cell clusters. Size of dots indicates the % of cells within each cluster that express the indicated gene. Colour of dots indicates the scaled expression of indicated gene across the clusters. Horizontal yellow bars at the top of the dot plot indicate which genes are expected to be increased and which are expected to be decreased in $T_{RM}$ cells[34]. For each cluster, genes/proteins which are significantly differentially expressed in the direction expected for $T_{RM}$ cells are marked with an asterisk. Vertical yellow bar indicates clusters meeting criteria to be defined as “potential $T_{RM}$ cell clusters”. Differential expression of SCTransformed RNA for each cluster compared to all other cells was calculated using the Wilcoxon signed rank test using the FindAllMarkers() function in Seurat. Adjusted p value <0.05 considered significant. **b)** Results (normalised enrichment score and FDR q-values) of GSEA indicating positive enrichment of human lung $T_{RM}$ cell signature[34] in clusters 1, 2, 4, 8, 14 and 15 when compared to pooled cells from non-potential-$T_{RM}$ clusters (clusters 3, 5, 6, 7, 9, 10, 11, 12, 13, 16, 17 and 18). Cluster 0 was not enriched for any of the tested $T_{RM}$ signatures (human lung and spleen CD4+ and CD8+ $T_{RM}$ signatures[34], normalised enrichment score visualised for cluster 0 is for human lung CD8+ $T_{RM}$ signature. (n=6 patients). **c)** Results (normalised enrichment score and FDR q-values) of GSEA analysis for each $T_{RM}$ cluster compared to pooled cells from other $T_{RM}$ clusters. Cluster 1 was not positively enriched for any of the tested gene sets compared to other $T_{RM}$ clusters therefore is not depicted here. However, cluster 1 had significant upregulation of *RORA* and *CCR6* and CD161 protein and therefore was classified as having a type-17 phenotype.

### Fig. S7 Durham et al.

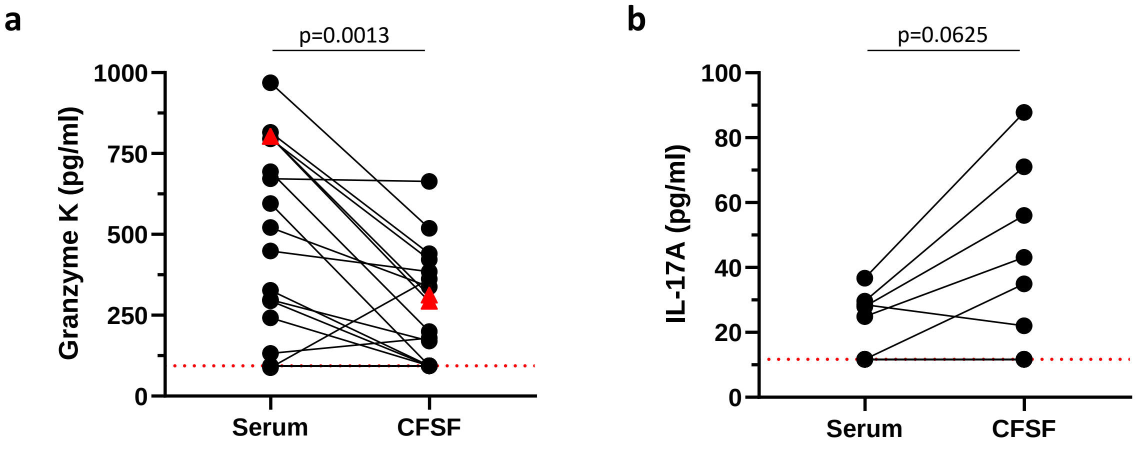


**Figure S7 Presence of granzyme K and IL-17A in serum and cell free synovial fluid in PsA**
**a,b)** Levels of **(a)** granzyme K (n=17) and **(b)** IL-17A (n=10) in paired serum and cell free SF (CFSF) from patients with PsA measured by ELISA. Red triangles indicate granzyme K in CFSF obtained from left and right knees from the same patient. Wilcoxon matched pairs signed rank test (two-tailed). Red dashed line indicated the lowest detection limit. Values below the lowest detection limit (n=5 serum, n=4 CFSF) were set to this value.

### Fig. S8 Durham et al.

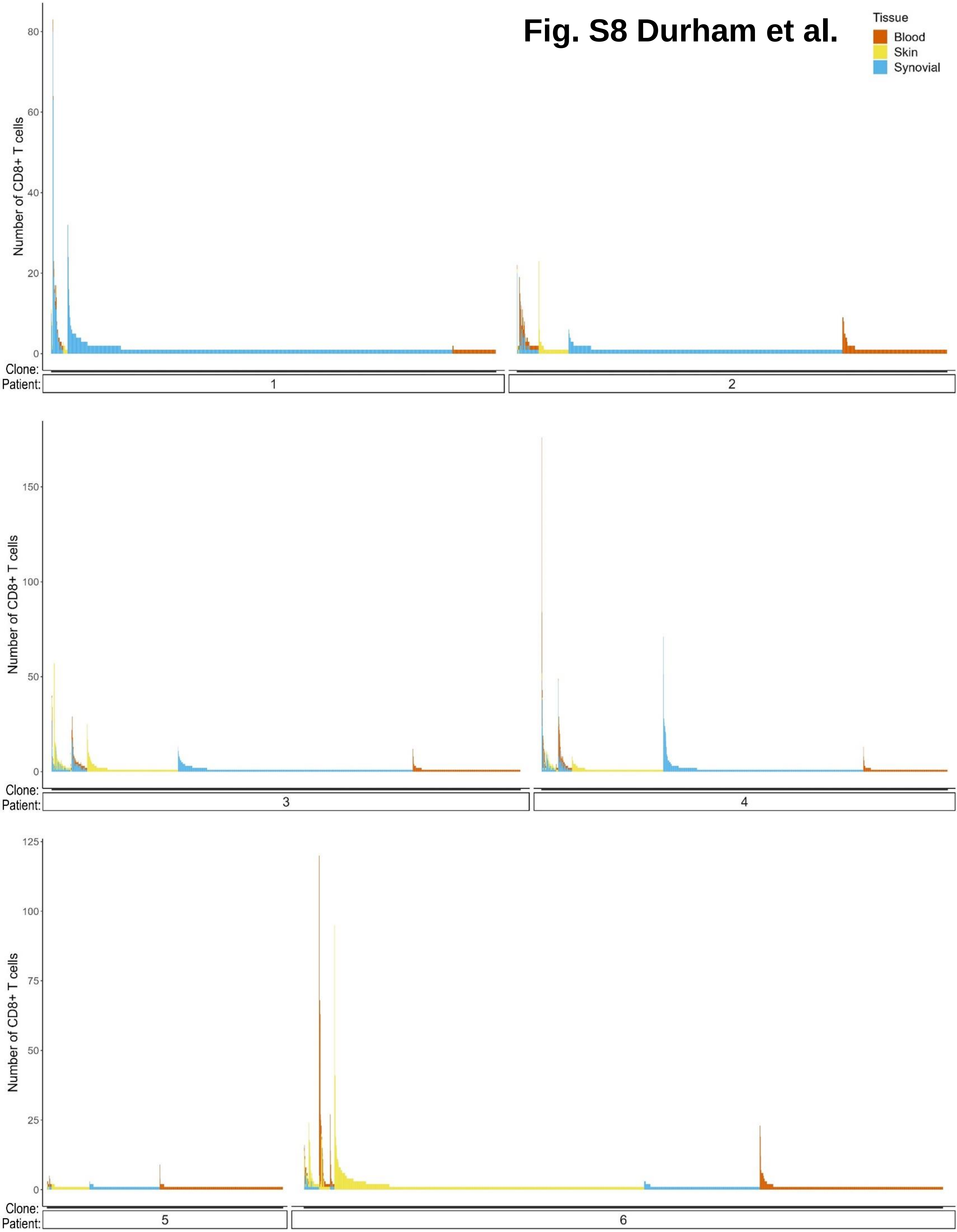

**Figure S8: Tissue location of all CD8+ T cell clones**

Bar chart visualising the size and tissue location of all CD8+ T cell clones. Each vertical bar represents a CD8+ T cell clone that is shared between the skin and the joint. The height of the bar represents the number of CD8+ T-cells within that clone and the colour represents the tissue that the cells were detected in (red = blood, yellow = skin, blue = joint). Clones are split by patient. Within each patient clones are ordered into triple shared clones, followed by dual shared clones (clones that were detected in the skin and the joint, in the joint and blood and in the skin and blood) followed by clones that were only detected in a single tissue (skin, joint or blood).

### Fig. S9 Durham et al.

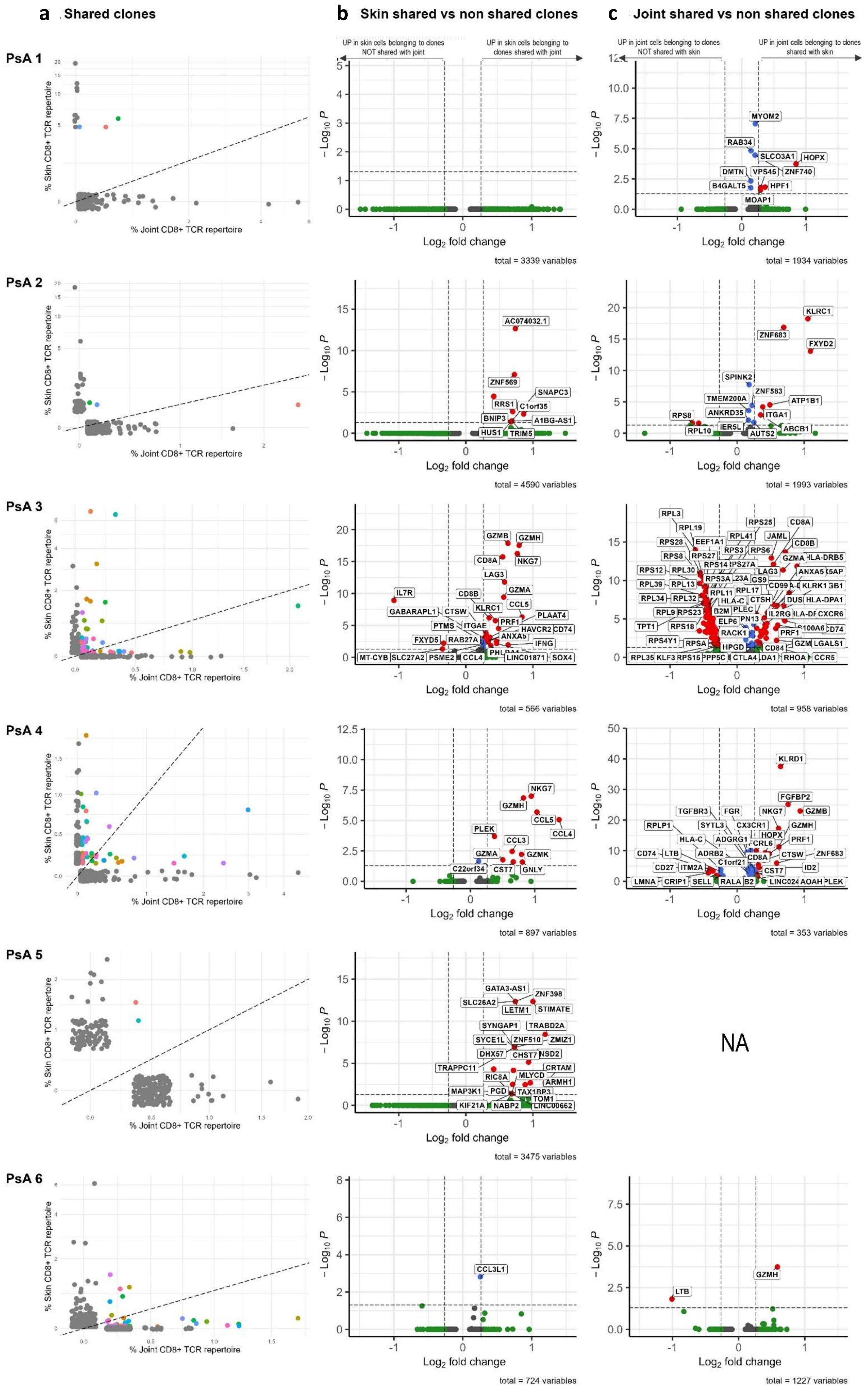

**Figure S9: CD8+ clones shared between skin and synovium**

**a)** Scatterplots for each patient showing the frequency of CD8+ T cell clones in skin and joint. Each dot represents a CD8+ T cell clone and the position of the dot along the x and y axes indicates the frequency of that clone in the joint and skin respectively. Scatterplot for PsA 3 is duplicated from Figure 3A for clarity. **b)** Volcano plot showing significantly differentially expressed genes between skin cells expressing TCRs which are shared with joint (i.e. shared clones) vs. skin cells expressing TCRs which are not shared with joint (i.e. non-shared clones) for each individual patient. Differential genes calculated using FindMarkers() in Seurat. **c)** Volcano plot showing differentially expressed genes between joint shared clones vs. joint non shared clones for each individual patient. NB: Differentially expressed genes not calculated for PsA 5 joint due to insufficient number of cells belonging to shared clones in the joint ($n < 3$).

**Fig. S10 Durham et al.**

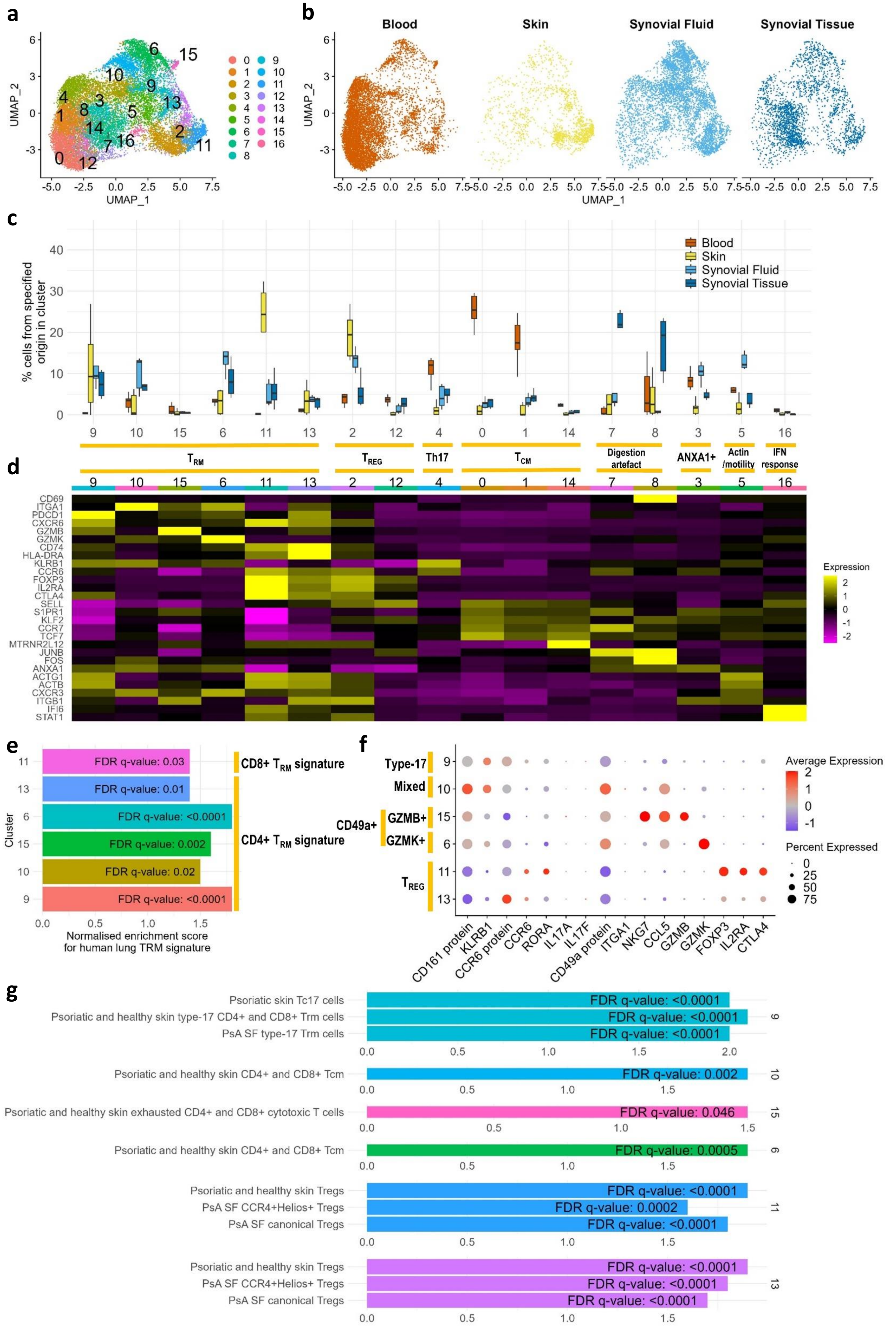

**Figure S10: Integrated analysis of 17,700 memory CD4+ T cells from paired samples of blood, lesional skin epidermis and ST and/or SF from inflamed knees from 6 patients with PsA (n=6 patients).**

**a)** UMAP with cells coloured according to the 17 cell populations obtained after Seurat clustering. **b)** UMAPs split by tissue of origin. **c)** Boxplot visualising the distribution of CD4+ T cells from each tissue across the 17 clusters. Clusters are grouped into groups with similar phenotypes. **d)** Heatmap visualising expression of specified genes across the clusters. **e)** Results (normalised enrichment score and FDR q-values) of gene set enrichment analysis showing positive enrichment of human lung $T_{RM}$ cell signature[34] in clusters 6, 9, 10, 11, 13 and 15 when compared to pooled cells from non-potential-$T_{RM}$ clusters. **f)** Dot plot depicting expression of select genes across the CD4+ $T_{RM}$ clusters. Size of dots indicates the % of cells within each cluster that express the indicated gene. Colour of dots indicates the scaled expression of indicated gene across all of the CD4+ $T_{RM}$ clusters. Yellow bars indicate clusters with similar phenotypes. **g)** Results of gene set enrichment analysis for each $T_{RM}$ cluster compared to pooled cells from other $T_{RM}$ clusters.

Fig. S11 Durham et al.

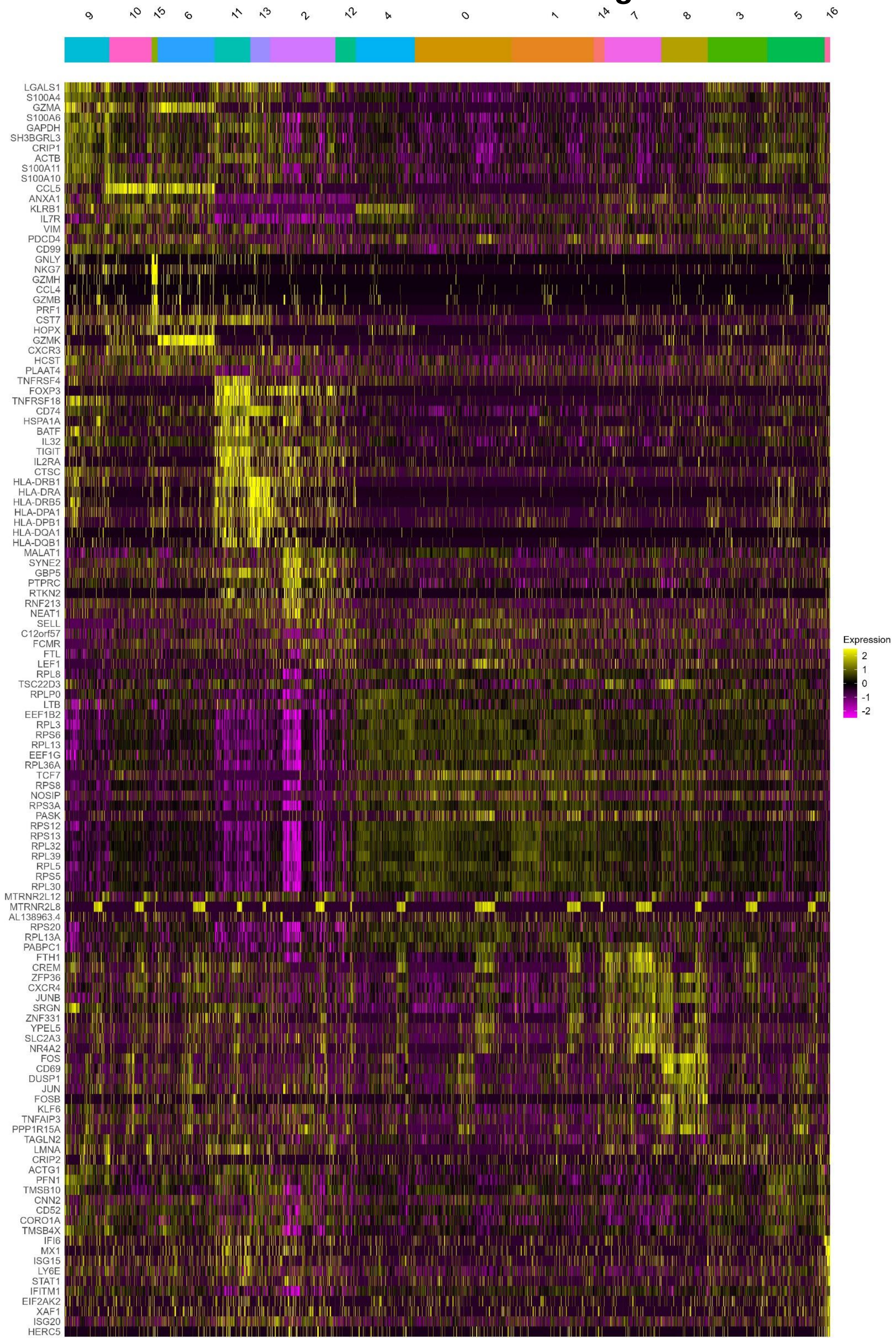

**Figure S11: Heatmap visualising the top 10 genes differentially expressed by each of the 17 CD4+ T cell clusters in the CD4+ T cell analysis.** Clusters grouped into groups with similar phenotypes. Differential expression by SCTransformed RNA was calculated using the Wilcoxon signed rank test using the FindConservedMarkers() function Seurat. FindConservedMarkers() mitigates for potential batch effect between patients by performing differential gene expression testing for each patient separately and combining the p-values using meta-analysis methods from the MetaDE R package. Combined p value $< 0.05$ was used to identify significantly differentially expressed genes. Log2FC for each of the 6 patients was then averaged and genes were then ranked in order of average Log2FC change.

**Supplementary Table 1: Antibodies used for Fluorescence Activated Cell Sorting and CITE-seq**

| **Fluorescence – conjugated antibodies** | | |
|---|---|---|
| **Antibody** | **Clone** | **Manufacturer** |
| CD14 – APCcy7 | REA599 | Miltenyi |
| CD3 – PEcy7 | UCHT1 | Biolegend |
| CD45 – BUV395 | HI30 | Becton Dickinson UK Ltd |
| CD27 – APC | O323 | Biolegend |
| CD45RA – BV711 | HI100 | Biolegend |
| **TotalSeqC antibodies** | | |
| **Antibody** | **Clone** | **Manufacturer** |
| PD1 | EH12.2H7 | Biolegend |
| CCR6 | G034E3 | Biolegend |
| CD103 | Ber-ACT8 | Biolegend |
| CD69 | FN50 | Biolegend |
| CD161 | HP-3G10 | Biolegend |
| CD49a | TS2/7 | Biolegend |
| CXCR6 | K041E5 | Biolegend |
| CD8A | RPA-T8 | Biolegend |
| Anti-human Hastags | LNH-94; 2M2 | Biolegend |

**Supplementary Table 2: Hashtags used to stain each patient sample**

Biolegend TotalSeq-C hashtags (anti-human, clone LNH-94 & 2M2, isotype: mouse IgG1)

| **Patient** | **Hashtag** | | | | | | |
|---|---|---|---|---|---|---|---|
| | **Blood** | | **SF** | | **ST** | | **Skin** |
| PsA 1 | 1 | 2 | 3 | 4 | 6 | | 5 |
| PsA 2 | 1 | 2 | 3 | 4 | 6 | | 5 |
| PsA 3 | 1 | 2 | 3 | 4 | 6 | | 5 |
| PsA 4 | 1 | 2 | 3 | | 4 | 5 | 6 |
| PsA 5 | 1 | 2 | NA | | 3 | 4 | 5 |
| PsA 6 | 1 | 2 | 3 | | 6 | | 5 |
